# An NADH-controlled gatekeeper of ATP synthase

**DOI:** 10.1101/2024.08.22.609182

**Authors:** Fabian Schildhauer, Petra SJ Ryl, Simon M Lauer, Swantje Lenz, Ayşe Berçin Barlas, Vasileios R Ouzounidis, Kate Jeffrey, Daniel-Cosmin Marcu, Francis J O’Reilly, Andrea Graziadei, Marchel Stuiver, Kita Schmidt, Helge Ewers, Christian MT Spahn, Ezgi Karaca, Karl Emanuel Busch, Dhanya Cheerambathur, David Schwefel, Juri Rappsilber

## Abstract

ATP fuels crucial cellular processes and is obtained mostly by oxidative phosphorylation (OXPHOS) at the inner mitochondrial membrane. While significant progress has been made in mechanistic understanding of ATP production, critical aspects surrounding its substrate supply logistics are poorly understood. We identify an interaction between mitochondrial apoptosis-inducing factor 1 (AIFM1) and adenylate kinase 2 (AK2) as gatekeeper of ATP synthase. This interaction is NADH-dependent and influenced by glycolysis, linking it to the cell’s metabolic state. Genetic interference with AIFM1/AK2 association impedes the ability of *Caenorhabditis elegans* animals to handle altered metabolic rates and nutrient availability. Together, the results imply AIFM1 as a cellular NADH sensor, placing AK2 next to the OXPHOS complexes for local ADP regeneration as the substrate for ATP synthesis. This metabolic signal relay balances ATP synthase substrate supply against ATP conservation, enabling cells to adapt to fluctuating energy availability, with possible implications for AIFM1-related mitochondrial diseases.

**Highlights:**

- Discovery of AIFM1/AK2 interaction as a gatekeeper of mitochondrial ATP synthase
- AIFM1/AK2 interaction is NADH-dependent and linked to the cell’s metabolic state
- Disrupting AIFM1/AK2 impairs metabolic adaptation in *Caenorhabditis elegans*
- AIFM1 as NADH sensor, influencing ATP synthesis with mitochondrial disease implications

## Introduction

ATP is the central energy carrier of the cell, and its production is fundamental to life^1^. In aerobic organisms, the majority of ATP is obtained via the IM OXPHOS machinery, comprising five membrane protein complexes (CI-CV). Complexes CI-CIV form an electron transport chain (ETC) that converts mitochondrial NADH to an electrochemical gradient, by proton translocation into the intermembrane space (IMS). This then drives ATP synthesis by CV, the ATP synthase complex. Tight control of ATP synthesis is essential to maintain energy homeostasis, redox balance, integration with cellular metabolism, adaptation to environmental conditions, mitochondrial quality control, and metabolic flexibility. Accordingly, a variety of regulatory mechanisms evolved to tune OXPHOS and ATP synthase activity^2–7^. Unsurprisingly, defects in these critical mitochondrial pathways are linked to diseases and ageing^8–13^.

An important regulatory aspect of any catalyst is the control of substrate supply and product removal. Indeed, the rate increase of ATP synthesis in muscles during exercise is driven by available ADP and P_i_ concentrations^14^. Efficient ATP extrusion into the cytoplasm is critical to avoid negative feedback from high ATP concentrations to the Krebs cycle and the ETC, meant to guard cells against OXPHOS overactivity^15–18^. Accordingly, an array of specific transport proteins evolved such as the mitochondrial phosphate carrier (PHC), magnesium transporter (MRS2), and the adenine nucleotide translocases (ANT1-4) to transport ATP synthesis substrates and extrude the reaction product across the impermeable IM^19–21^. Moreover, AK2, a soluble protein in the IMS, catalyses the reversible reaction ATP + AMP ↔ 2 ADP, thereby maintaining the local balance of adenine nucleotides and ensuring an adequate supply of ADP for ATP synthase^22–24^. The physiological importance of AK2 is emphasised by AK2 deficiency leading to OXPHOS defects, impairing haematopoiesis, and causing reticular dysgenesis, a form of human severe combined immunodeficiency^25–27^.

AIFM1 is an IMS flavoprotein tethered to the IM by an N-terminal transmembrane segment, initially described to be released from mitochondria upon apoptosis induction, affecting chromatin condensation and DNA fragmentation in a caspase-independent manner^28,29^. More recently, critical functions of AIFM1 in mitochondrial bioenergetics emerged. AIFM1 acts as an IMS import receptor to recruit and stabilise the MIA40/CHCHD4 oxidoreductase, mediating oxidative folding and import of several IMS and IM proteins, among them CI, CIII, CIV subunits and assembly factors, and interestingly also AK2^30–34^. Accordingly, AIFM1 deficiency compromises OXPHOS function^35,36^, and mutations in AIFM1 cause mitochondrial disease, including Cowchock syndrome, mitochondrial encephalopathy, combined OXPHOS deficiency, and deafness with peripheral neuropathy^37^.

OXPHOS complexes represent one of the best-understood biological systems in terms of biochemistry, structure and (patho)physiology^2,3,10–13,38–42^. However, the regulation of substrate supply deserves further investigation. Here, we apply chemical crosslinking and mass spectrometry, in conjunction with biochemical, structural, and cellular analyses, to demonstrate the NADH-dependent interaction between AIFM1 and AK2, placing them in spatial proximity to OXPHOS complexes III-V. The findings reveal a mechanistic connection between IMS NADH levels and the ATP synthesis substrate supply to optimise OXPHOS function. Moreover, these insights may contribute to a better understanding of the functional implications of disease-associated mutations in AIFM1.

## Results

### Crosslinking mass spectrometry interactome of isolated mitochondria

To reveal mitochondrial protein-protein interactions (PPIs) inaccessible to conventional identification approaches prone to cell lysis artefacts, and to gain insights into PPI topology, mitochondria isolated from human K-562 cells were subjected to *in situ* crosslinking using the membrane permeable reagent disuccinimidyl sulfoxide (DSSO). Subsequently, the crosslinked preparation was lysed, followed by proteolytic digestion, two-dimensional peptide ion exchange chromatography, and liquid chromatography (LC)-mass spectrometry (MS) data acquisition. Of the total detected protein intensity in the soluble and insoluble fraction, respectively, 72% and 40% arose from annotated mitochondrial proteins (Fig. S1A). We identified 7,033 crosslinks within single proteins (self-links) and 1,444 heteromeric crosslinks with a 5% residue-pair false discovery rate (FDR), representing 501 distinct PPIs at a 2% PPI-FDR. Using the example of complex IV, one can see that the crosslink data not only reveal putative interaction partners but extend to residue-resolved interaction footprints (Fig. 1A). We then focused further analysis on mitochondrial proteins and their crosslinking network (Fig. S1B). Of these, the mitochondrial matrix (MM) and the IM were the most abundant submitochondrial compartments, with fewer proteins identified from the IMS or outer membrane (OM), aligning with the overrepresentation of MM and IM proteins in the mitochondrial proteome^43^. The network’s largest fraction of co-purified non-mitochondrial proteins originated from the endoplasmic reticulum and plasma membrane/endosomal compartments, mostly representing crosslinks between mitochondrial proteins and chaperones or sorting factors. Importantly, our analysis yielded a highly interconnected and topologically well-separated crosslink network representing physiological meaningful PPIs according to STRING database annotation. Moreover, the fraction of crosslinks falling into available structures supports the resolution of crosslinking in the range of 3 nm for proteins within their cellular context (Fig. S1C-G).

**Fig. 1:**
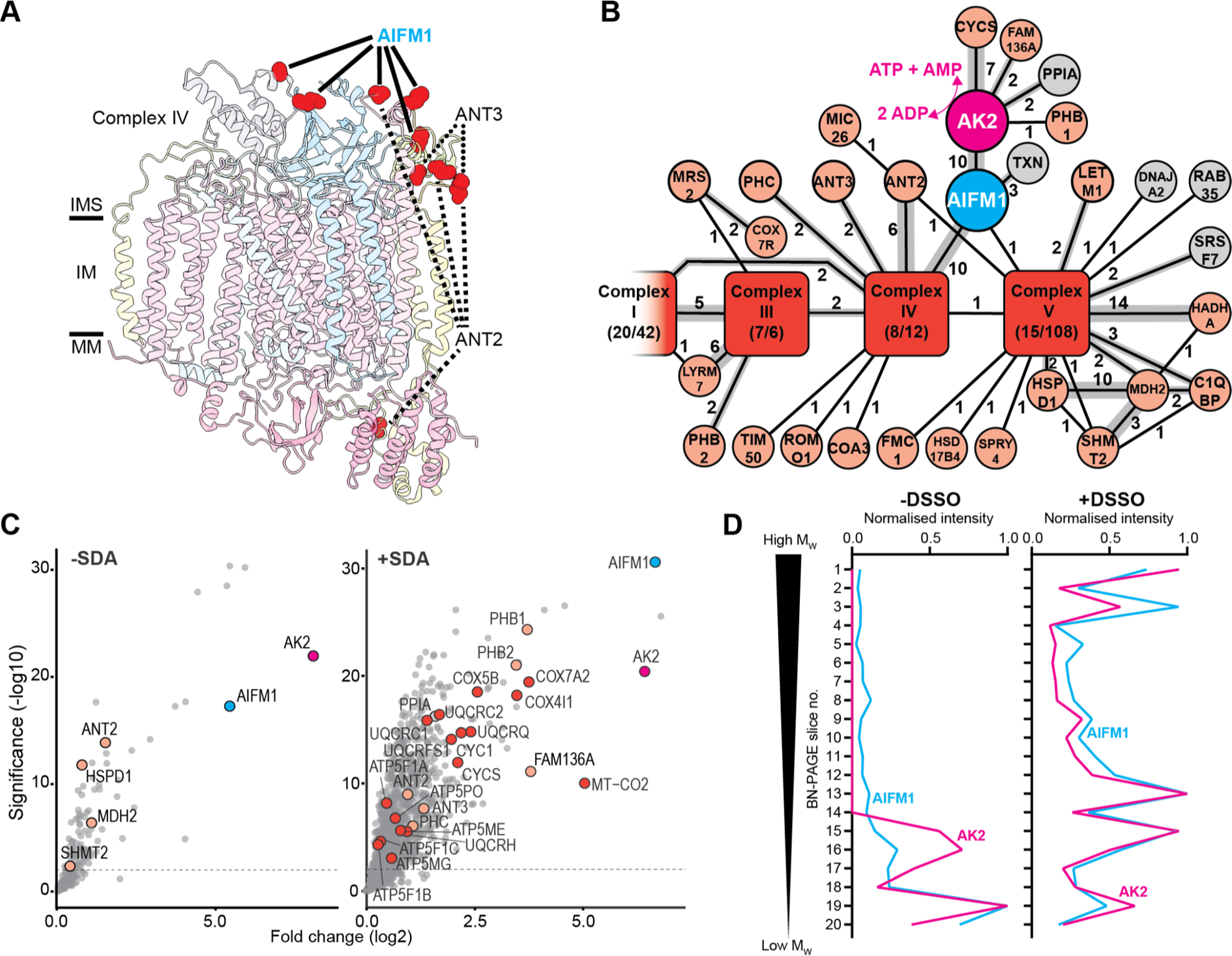
An AIFM1/AK2 complex in close vicinity to OXPHOS assemblies. (A) Mapping of DSSO crosslinks from the indicated proteins on the structure of human CIV (PDB 5z62^44^). (B) Sub-network of crosslinks around OXPHOS CIII-CV. AIFM1 is coloured blue, AK2 magenta, mitochondrial proteins light pink, and OXPHOS complexes red brown. Numbers beside the network edges indicate unique crosslinked residue pairs between the respective proteins. (C) AP-MS: proteins enriched in FLAG pull-downs of non-crosslinked or in-cell SDA (6 mM) UV-crosslinked, FLAG-AK2-transfected 293T cells, compared to non-crosslinked or SDA-crosslinked empty vector controls. Proteins from B are labelled and coloured accordingly. (D) Complexome profiling of isolated non-crosslinked (left) and crosslinked (0.25 mM DSSO, right) mitochondria. Relative MS label-free quantification (LFQ) intensities are plotted for AIFM1 and AK2 across all gel slices.

**Fig. S1:**
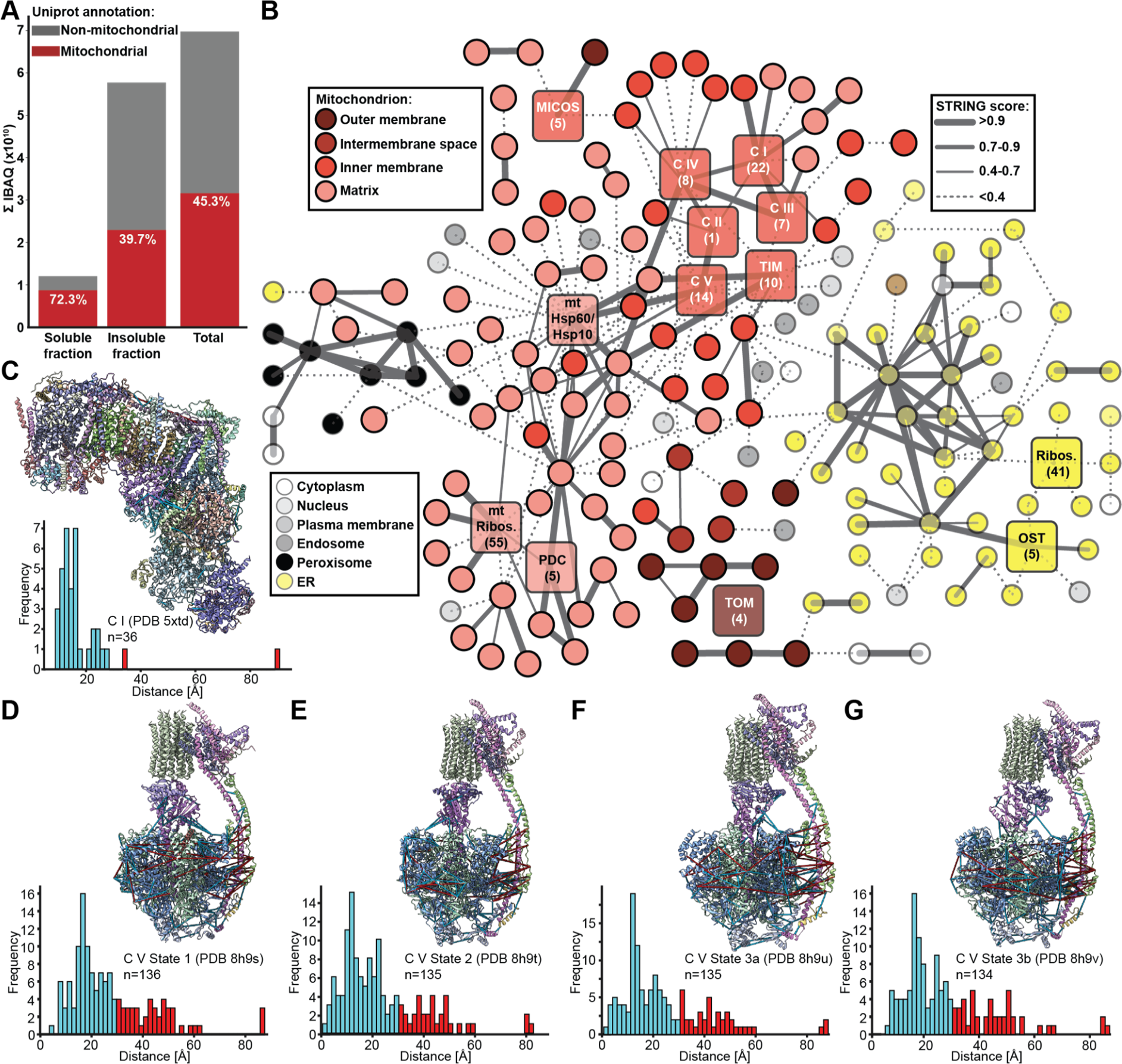
Detailed analysis of the mitochondrial DSSO crosslinking network, related to Fig. 1. (A) Intensity-based absolute quantification (IBAQ) of protein groups identified in the soluble and insoluble fraction of the mitochondrial preparation. The IBAQ contribution from proteins annotated as mitochondrial in the UniProt database is coloured red, with the respective percentage indicated. (B) Simplified view of the crosslinking network surrounding all identified mitochondrial proteins, with the round nodes representing individual proteins and the squares larger protein complexes, coloured according to their subcellular (from UniProt) or submitochondrial (from MitoCarta 3.0) localisation. The thickness of the network edges is proportional to the STRING score, which indicates the confidence for a functional association between the connected proteins, with values >0.4 indicating medium confidence, >0.7 high confidence, and >0.9 highest confidence. MICOS - mitochondrial contact site and cristae organizing system; mt - mitochondrial; OST - oligosaccharyltransferase; PDC - pyruvate dehydrogenase; TIM/TOM - translocase of the inner/outer mitochondrial membrane. (C-G) Mapping of crosslinks falling into available structures of CI (PDB 5xtd^45^) and CV (PDB 8h9s, 8h9t, 8h9u, 8h9v^46^). Crosslink distances are plotted in the histograms below. Crosslinks that are compatible with the maximal length imposed by the crosslinker chemistry are coloured blue, otherwise red.

**Fig. S2:**
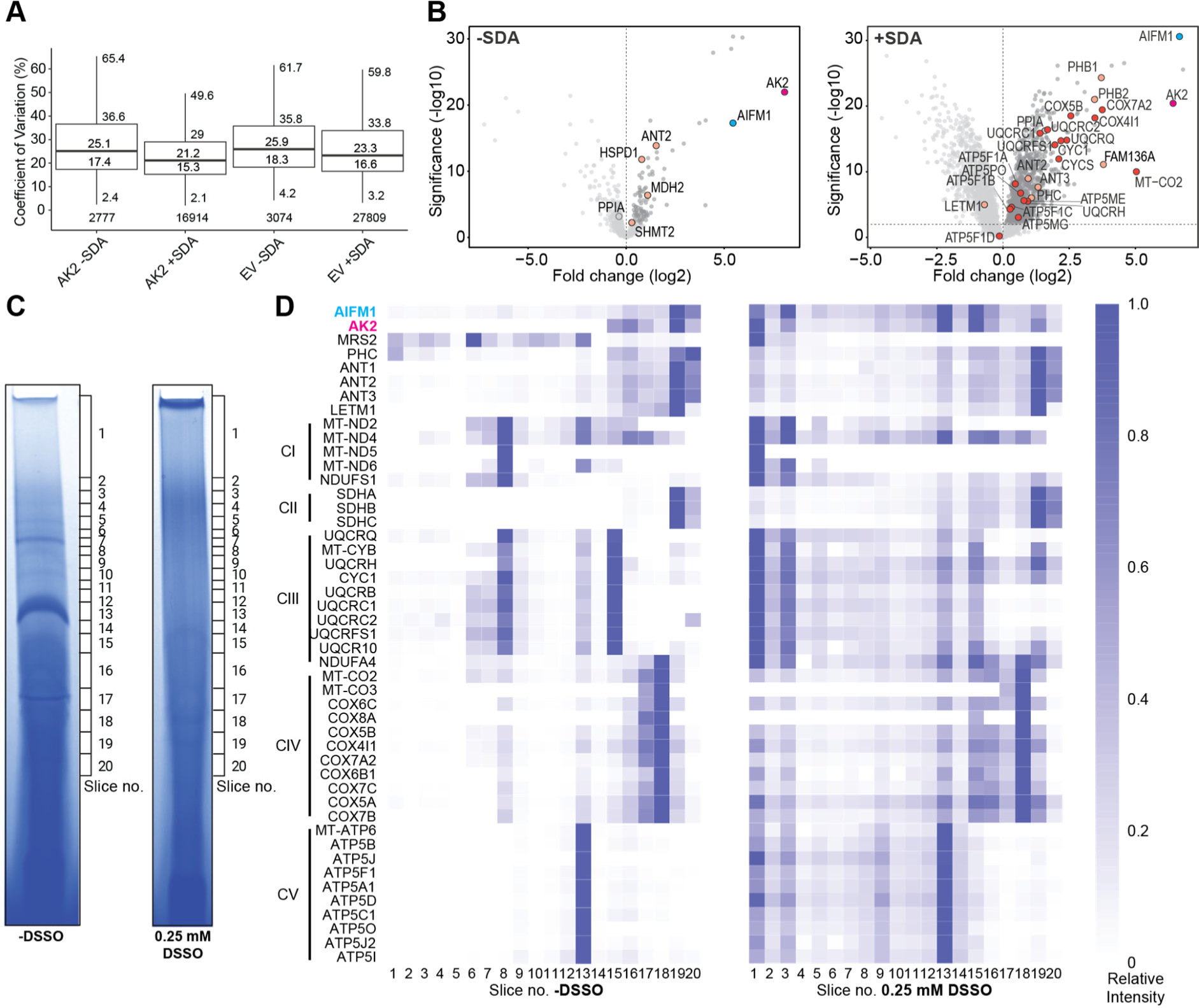
FLAG-AK2 AP-MS and complexome profiling of isolated mitochondria, related to Fig. 1. (A) AP-MS: peptide-level coefficients of variation indicating the reproducibility of replicate measurements, calculated by the MS-DAP platform^47^. Data were analysed as pools of biological triplicates, each one injected in technical triplicates into the mass spectrometer. (B) AP-MS: full volcano plots as in Fig. 1C but including negative fold change values. (C) Complexome profiling: coomassie blue-stained BN-PAGE gel lanes of isolated mitochondria untreated (left) or treated with 0.25 mM DSSO. Positions of the slices used for LC-MS analysis are indicated. (D) Complexome profiling: heatmap of relative LFQ intensities determined from BN-PAGE gel slices, normalised to the maximum intensity of each protein across all slices.

### An AIFM1/AK2 complex in close vicinity of the OXPHOS machinery

To obtain mechanistic insight into substrate supply logistics of ATP production, the analysis was directed towards a PPI subnetwork surrounding the terminal OXPHOS complexes CIII, CIV and ATP Synthase/CV. Prominently, we noticed several crosslinks connecting AK2 to AIFM1, which in turn was linked to CIV and CV (Fig. 1B).

To substantiate the physical interaction between AIFM1 and AK2, we conducted affinity purification (AP)-MS analyses of 293T cells transfected with epitope-tagged AK2 (Fig. S2A). Quantitative comparison to control transfections confirmed the co-purification of AIFM1 and, to less extent, ANT2 (Fig. 1C, left panel, Fig. S2B). Moreover, to verify the subcellular context of the AIFM1/AK2 complex, we performed similar experiments after in-cell photo-crosslinking using the heterobifunctional, UV-activatable reagent succinimidyl 4,4’-azipentanoate (SDA). These data showed an accumulation of OXPHOS subunits (CIV, CIII, CV, in the order of enrichment) and several mitochondrial proteins identified from the DSSO crosslinking network shown in Fig. 1B, among them all direct AK2 interactors, and ANT2/3 (Fig. 1C, right panel, Fig. S2B).

Additionally, complexome profiling of isolated mitochondria was performed, in the absence and presence of DSSO. Protein assemblies were separated by size through blue native gel electrophoresis (BN-PAGE) and migration profiles were determined by gel slicing and LC-MS (Fig. S2C). Without crosslinking reagent, AIFM1 and AK2 co-migrated in lower molecular weight fraction 19 (Fig. 1D, left panel, Fig. S2D). The addition of DSSO led to a shift of the co-peaks of AK2 and AIFM1 to higher molecular weight fractions 15, 13, and 3, which also contained ANT1-3 and, to varying degrees, OXPHOS CIII and CIV subunits (fraction 15), CV subunits (fraction 13), or CI, CIII, CIV, and CV subunits (fraction 3) (Fig. 1D, right panel, Fig. S2D).

Altogether, these results are in line with our large-scale DSSO crosslinking network analysis, supporting the existence of an IMS AIFM1/AK2 protein complex adjacent to ANTs and the IM OXPHOS assemblies CIII-CV.

### The AIFM1/AK2 interaction is NADH-dependent

Intrigued by the possible role of AIFM1 in inducing proximity between AK2 and sites of OXPHOS/ATP synthesis, we set out to characterise the AIFM1/AK2 interaction in more detail. AlphaFold multimer structure prediction yielded a high-confidence model, where the C-terminus of AK2 (residues 232-239) was inserted as a β-strand into the AIFM1 C-terminal (C-) domain, leading to a stacked β-sheet structure (Fig. S3).

**Fig. S3:**
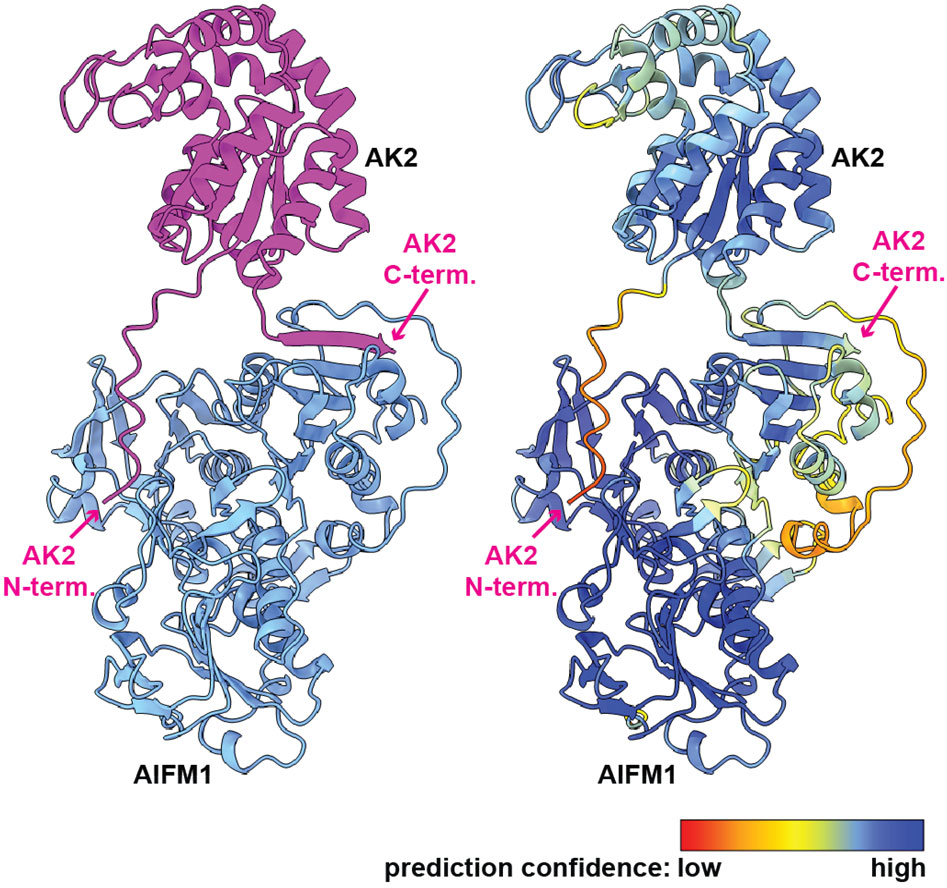
AlphaFold multimer structure prediction of the AIFM1/AK2 complex, related to Fig. 2. The amino acid sequences of human AIFM1 (UniProt ID O95831) and human AK2 (UniProt ID P54819) were used as an input for AlphaFold v2.3.2 structure prediction^48^. The resulting model is shown in cartoon representation, in the left panel coloured according to protein type (AIFM1 - blue, AK2 - pink) and in the right panel according to the residue-based confidence score pLDDT, ranging from 0 (red) to 100 (dark blue).

To ascertain the *in silico* model, recombinant human AIFM1 lacking its N-terminal signal sequence and transmembrane segment (AIFM1 Δ1-102, from here on AIFM1), and AK2 were recombinantly produced and purified. Size-exclusion chromatography (SEC) and mass photometry revealed no stable AIFM1/AK2 interaction in the absence of cofactors (Fig. 2A, B, pink, blue and grey traces). The addition of NADH to AIFM1 triggered dimer formation, as described previously^49–51^ (Fig. 2A, B, green traces), and elicited SEC co-elution of approximately 50% of the equimolarly added AK2 together with AIFM1 (Fig. 2A, orange trace). These data demonstrated stable, NADH-dependent AIFM1/AK2 interaction, with a 2:1 stoichiometry (AIMF1_2_:AK2) confirmed by mass photometry (Fig. 2B, orange traces).

**Fig. 2:**
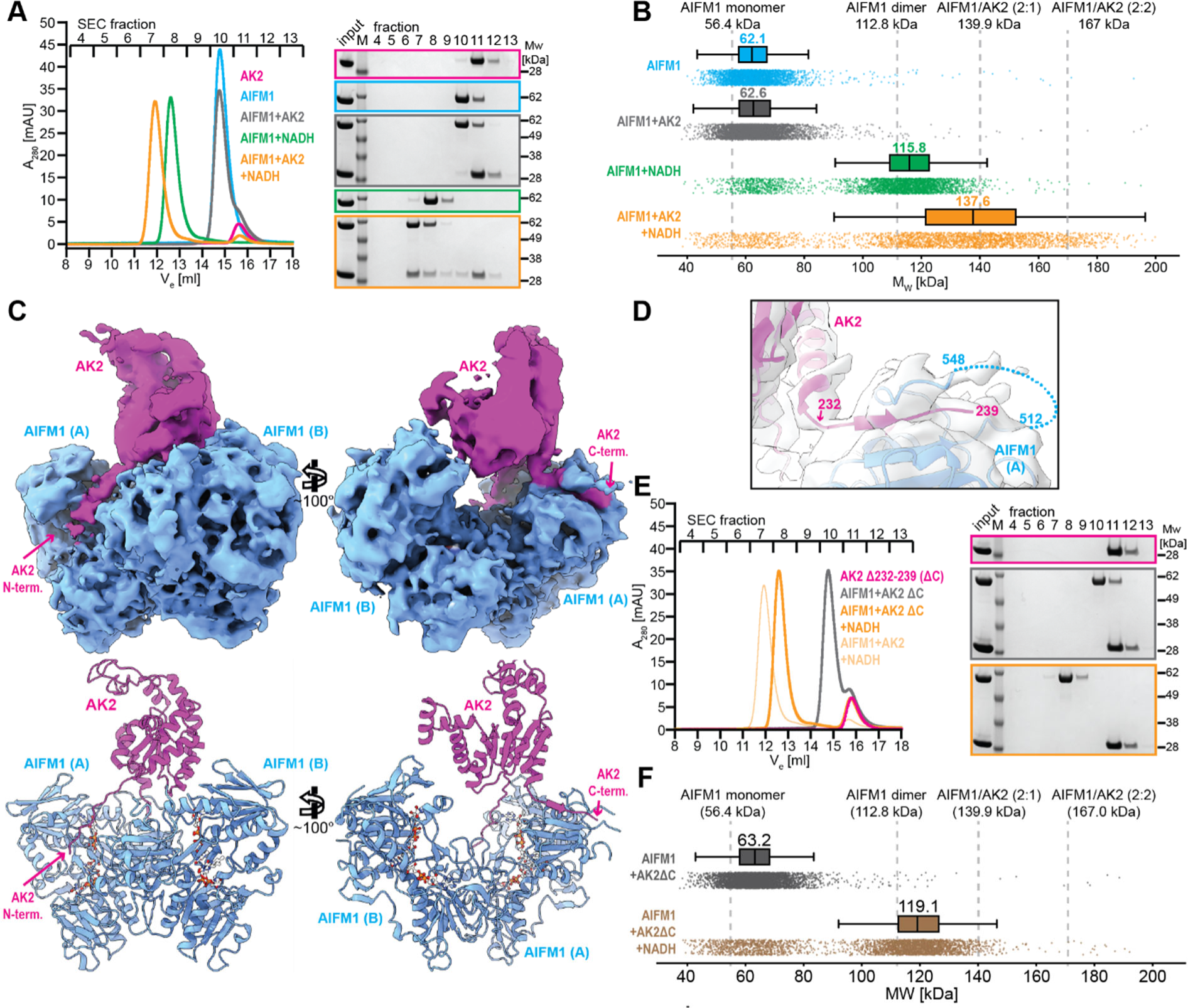
AIFM1/AK2 interaction depends on NADH and the AK2 C-terminus. (A) *In vitro* reconstitution of AIFM1, AK2 and complexes, followed by analytical SEC, in the absence and presence of NADH. SDS-PAGE analyses of SEC fractions are shown next to the chromatogram. (B) Mass photometry analyses of AIFM1 and complexes with AK2 in the absence and presence of NADH. Box-and-whisker plots of apparent molecular weights are shown above the raw data counts, with the median molecular weights indicated (for the range 40-90 kDa in the absence of NADH or 90-200 kDa in the presence of NADH). Theoretical molecular weights of AIFM1 and AIFM1/AK2 assemblies are shown on top of the graph and indicated by the grey dashed lines. (C) Cryo-EM structure analysis of the AIFM1/AK2 complex. The upper panel shows two views of segmented cryo-EM density for AIFM1 (blue) and AK2 (magenta). The lower panel depicts the corresponding fitted molecular model. (D) Detailed view of the cryo-EM density around the AK2 C-terminus binding site. (E) *In vitro* reconstitution, followed by SEC, of AIFM1, an AK2 variant lacking the C-terminal residues 232-239 (AK2 ΔC), and an equimolar mixture of both proteins, in the absence and presence of NADH. The SEC trace of the AIFM1/AK2 WT complex from A is shown in pale orange for comparison. (F) Mass photometry analyses of AIFM1 incubations with AK2 ΔC in the absence and presence of NADH. Box-and-whisker plots of apparent molecular weights are shown above the raw data counts, with the median molecular weights indicated (for the range 40-90 kDa in the absence of NADH or 90-200 kDa in the presence of NADH). Theoretical molecular weights of AIFM1 and AIFM1/AK2 assemblies are shown on top of the graph and indicated by the grey dashed lines.

**Fig. S4:**
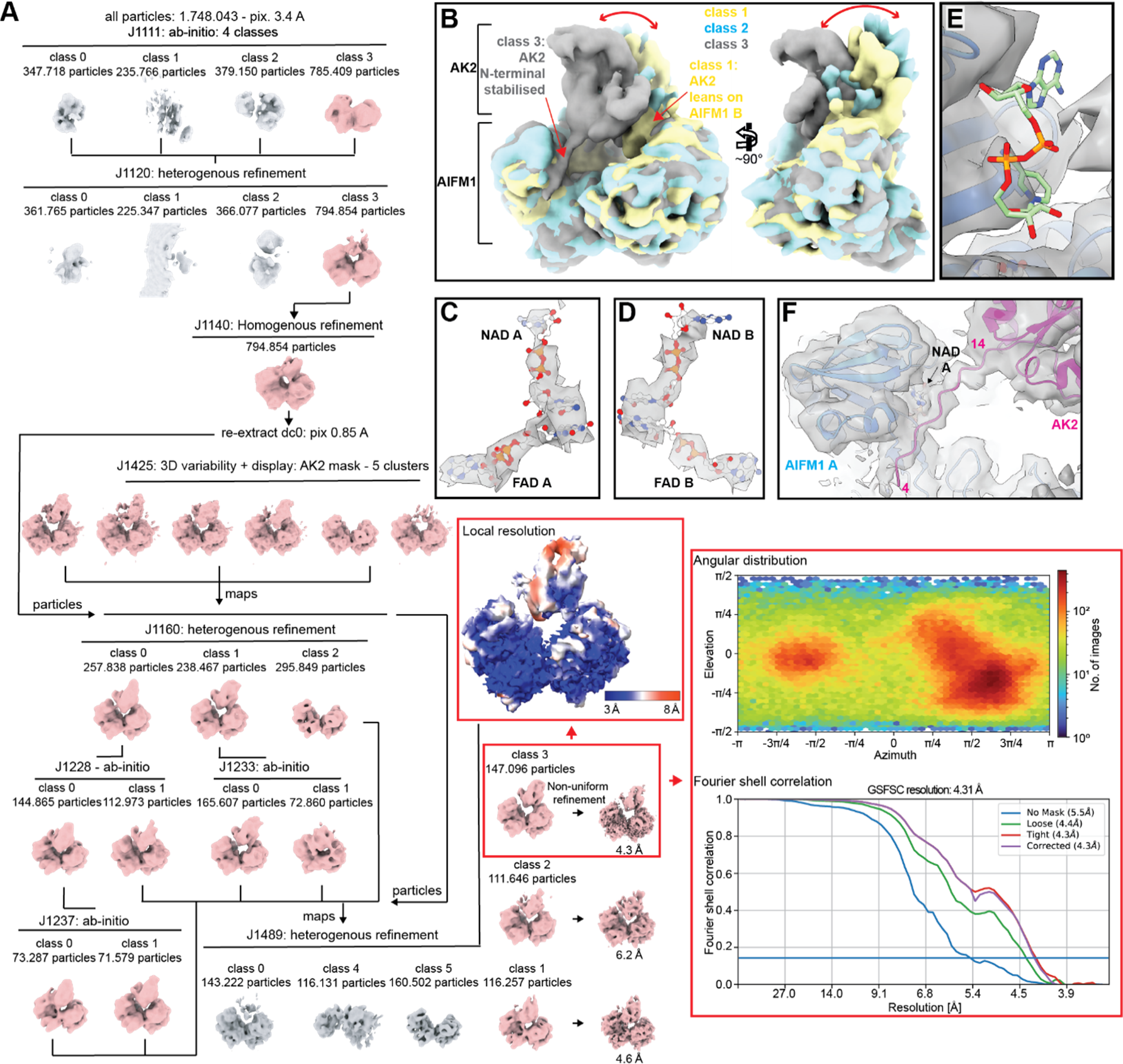
Cryo-EM single particle analysis of the AIFM1/AK2 complex, related to Fig. 2. (A) Cryo-EM data processing flowchart. All procedures have been performed using the program CryoSparc^50,52^. (B) Superposition of 3D classes 1-3 obtained from the last heterogeneous refinement step (J1489). Density portions corresponding to the AIFM1 homodimer have been aligned, to visualise the variations in AK2 positioning relative to AIFM1 (highlighted by the red double arrows). Contacts between AK2 and AIFM1, observable in classes 1 and 3, are indicated by arrows. (C) Class I cryo-EM density confirming the presence of the FAD/NADH charge transfer complex in AIFM1 protomer A. (D) Class I cryo-EM density confirming the presence of the FAD/NADH charge transfer complex in AIFM1 protomer B. (E) Lack of cryo-EM density in the secondary NADH binding site observed in a previous crystal structure^50^. (F) Class I cryo-EM density indicating the contact between the AK2 N-terminus and AIFM1.

### The AK2 C-terminus is the key determinant for its AIFM1-binding

Pursuing cryogenic electron microscopy (cryo-EM) to gain further mechanistic insights into the AIFM1/AK2 complexation revealed heterogeneity of AK2’s position relative to the AIFM1 homodimer, indicating a flexible tether between the two proteins (Fig. S4A, B). One 3D class, comprising 39% of all AK2-containing particles of the final classification round, was resolved to a global resolution of 4.3 Å (3-8 Å local resolution) and was used for fitting and refining molecular models of the AIFM1 homodimer and the AIFM1-bound AK2 AlphaFold prediction (Fig. 2C, Fig. S4A, Table S1). The cryo-EM density supported the presence of an FAD/NADH charge transfer complex consistent with the crystal structure of the NADH-bound human AIFM1 homodimer^50^ (Fig. S4C, D), although a secondary NADH binding site could not be confirmed (Fig. S4E). Altogether, the molecular model (i) rationalises the asymmetric 2:1 AIFM1:AK2 stoichiometry observed biochemically through the steric exclusion of a second AK2 molecule bound to the opposing AIFM1 protomer, in contrast to the expected 2:2 stoichiometry based on the symmetric arrangement of the AIFM1 dimer; (ii) validates the AlphaFold-based hypothesis of the AK2 C-terminus (residues 232-239) inserting as β-strand into the AIFM1 C-domain (Fig. 2D); and (iii) indicates contact between the extended AK N-terminus (residues 4-14) and the AIFM1 NADH-binding domain in at least a third of the AK2-containing particle population (Fig. 2C, Fig. S4F). The latter observation reflects the better resolution of this particular 3D class, as the additional stabilisation of AK2 by immobilisation of its N-terminus rigidifies the AIFM1/AK2 assembly. Strikingly, AK2 ΔC, lacking residues 232-239, did not stably interact with AIFM1 in the presence of NADH, identifying the C-terminus as the main AK2 determinant of AIFM1 interaction in these experimental settings (Fig. 2E, F).

### Allosteric mechanism of AK2 recruitment

To determine the molecular mechanism of AK2 recruitment, we compared our AK2/AIFM1 complex with available structures of oxidised monomeric and NADH-reduced dimeric AIFM1. NADH-binding triggers two allosteric pathways in AIFM1: one leading to dimerisation involving an interface in the FAD-binding domain, and the other causing the detachment of a C-terminal loop (C-loop, residues 509-559)^49–51^. In the oxidised form, most of the AIFM1 C-loop (residues 509-545) is structured through multiple hydrophobic and ionic interactions, including contacts with W196 and R201 within the Cβ-clasp motif connecting the FAD- and NADH-binding domains (Fig. 3A, upper panel). NADH-binding and subsequent charge transfer complex formation initiate allosteric Cβ-clasp remodelling and C-loop detachment (Fig. 3A, middle panel)^49^. This alleviates C-loop conformational constraints, allowing C-loop sections 509-512 and 546-559 to wrap around the AK2 C-terminus upon AK2 insertion (Fig. 3A, lower panel).

**Fig. 3:**
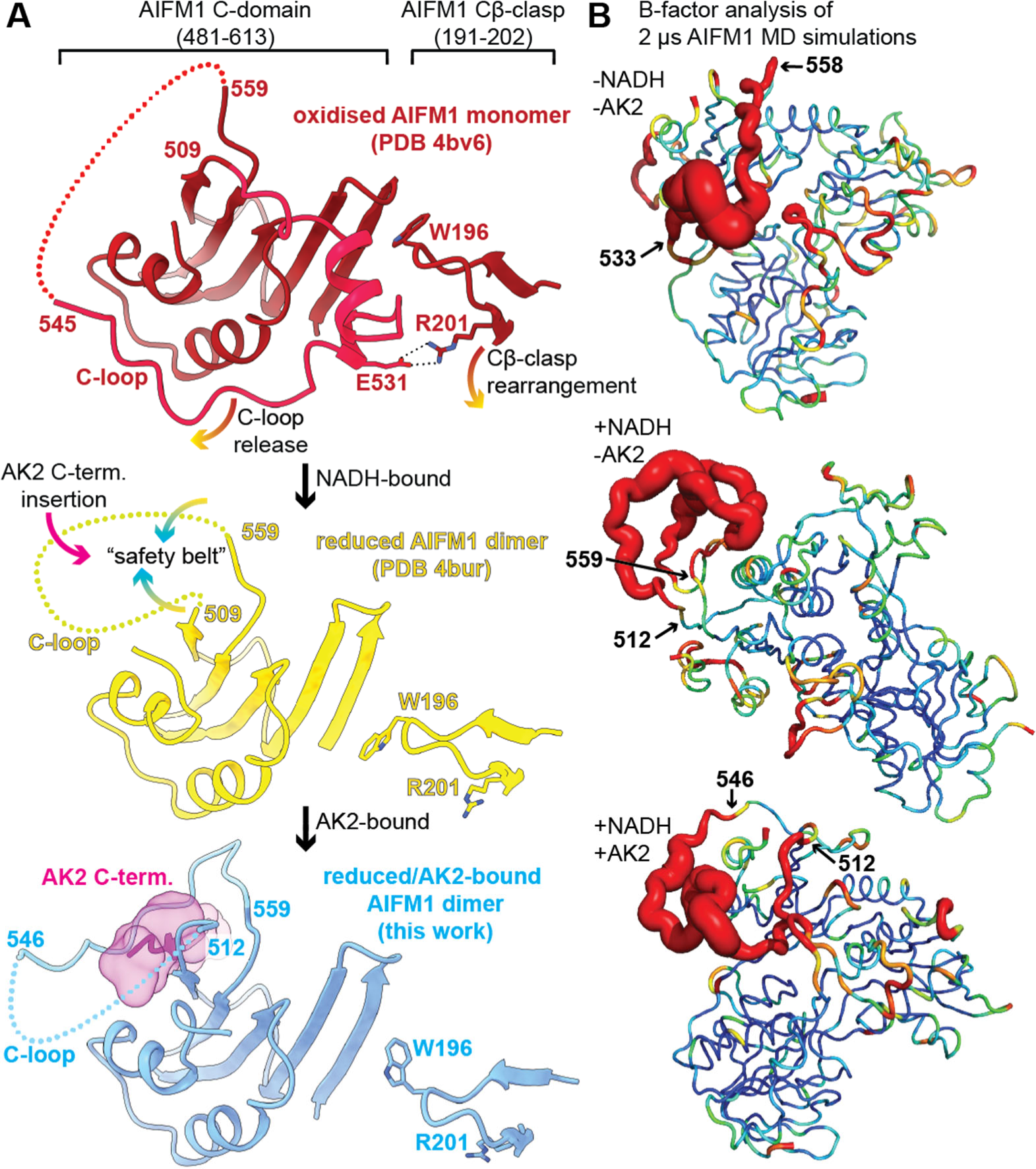
Mechanistic analysis of the NADH-dependent AIFM1/AK2 interaction. (A) Structural comparison of the C-domains of oxidised monomeric AIFM1^50^, NADH-reduced dimeric AIFM1, and AK2-bound, reduced dimeric AIFM1. Structural transitions between these states are indicated by the arrows. (B) B-factor flexibility analysis of the indicated MD simulations, mapped on the structure of AIFM1. The “-NADH/-AK2” analysis represents the AK2-unbound AIFM1 protomer from the AIFM1(FAD)_2_:AK2 trajectory (see Methods), the “+NADH/-AK2” represents the AK2-unbound protomer from the AIFM1(NAD+FAD)_2_:AK2 trajectory, and the “+NADH/+AK2” the opposing AK2-bound AIFM1 protomer from the same simulation.

To support these interpretations, we initiated all-atom molecular dynamics (MD) simulations. For this, we ran 4 µs (2x 2 µs) long MD simulations of AIFM1(NAD+FAD)_2_:AK2 and AIFM1(FAD)_2_:AK2 complexes (Fig. S5A, B). The simulations successfully replicated the flexible attachment of AK2 to AIFM1(FAD+NADH), as observed in the 3D classification of the cryo-EM data. Our trajectories converged to conformations involving contacts between the AK2 N-terminus and AIFM1, even though the simulations were started from an extended conformation of the AIFM1 C-loop (Supplemental Movie, Fig. S5A). Importantly, flexibility analyses of the resulting trajectories fully agreed with the above observations, demonstrating low mobility of the C-loop region 509-533 in the absence of NADH and AK2, high fluctuation of nearly all C-loop residues (512-559) upon NADH addition, and immobilisation of C-loop sections 509-512 and 546-559 upon further inclusion of AK2 in the simulation (Fig. 3B). Moreover, our analysis of MD trajectories showed that the allosteric modulation is driven by non-specific salt bridges formed between the C-loop and neighbouring charged AIFM1/AK2 amino acids (Fig. S5C). We observed a direct correlation between the number of these salt bridges and experimental structures: a higher number of salt bridges corresponds to an increase in the length of resolved regions. This finding highlights the crucial role of charged residues, which constitute approximately 25% of the C-loop segment. Taken together, the allosteric release frees the AIFM1 C-loop to fasten like a “safety belt” around the AK2 C-terminus, enabling stable AK2 association.

**Fig. S5.**
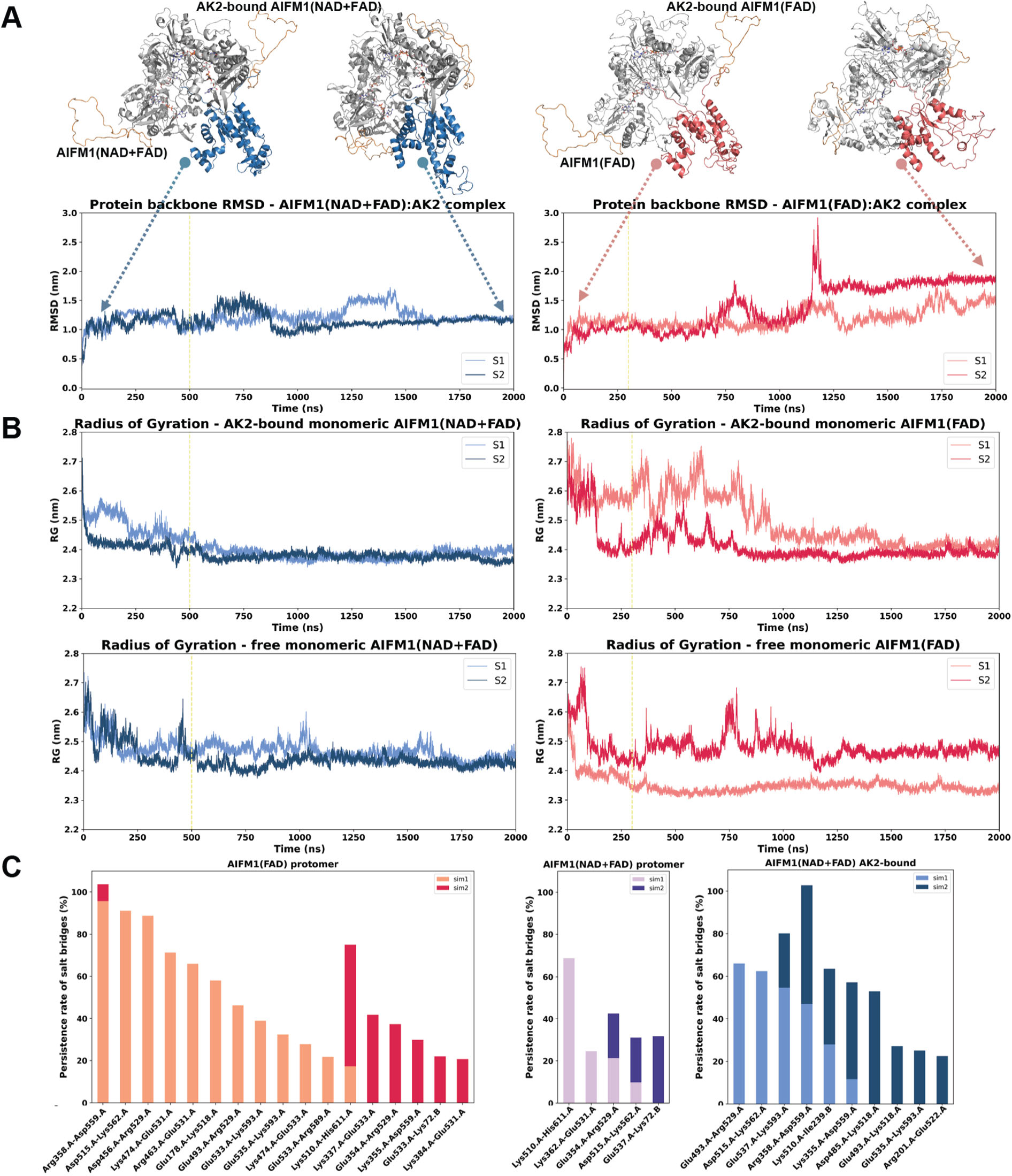
Quality control and detailed analysis of MD simulations of AIFM1(NAD+FAD)_2_:AK2 and AIFM1(FAD)_2_:AK2 complexes, related to Fig. 3. (A) Root Mean Square Deviation (RMSD) values calculated over backbone atoms of AIFM1(NAD+FAD)_2_:AK2 (left) and AIFM1(FAD)_2_:AK2 complexes during 2 µs long replica MD simulations (by taking the initial conformation as a reference where we did not impose any conformation on the flexible AIFM1 C-loop). The vertical lines indicate the time section considered as the system equilibration. The structural snapshots represent the initial and the final conformations of the complexes. Here, AK2s are depicted in blue in AIFM1(NAD+FAD):AK complex and salmon in AIFM1(FAD):AK2 complex. AIFM1 dimers are shown in gray and the C-loops are highlighted in orange. (B) Radius of gyration (Rg) evaluations of the of (i) AK2-bound monomeric AIFM1(NAD+FAD) (upper left) and AK2-unbound monomeric AIFM1(NAD+FAD) protomer (lower left), taken from AIFM1(NAD+FAD)_2_:AK2 simulations; (ii) AK2-bound monomeric AIFM1(FAD) (upper right) and AK2-unbound monomeric AIFM1(FAD) protomer (lower right), extracted from AIFM1(FAD)_2_:AK2 simulations. The vertical lines indicate the time section considered as the system equilibration. (C) Persistence rates of salt bridges formed by the AIFM1 C-loop in two replica simulations of protomer (monomeric) AIFM1(FAD), taken from the AIFM1(FAD)_2_:AK2 complex simulations, protomer (monomeric) taken from the AIFM1(NAD+FAD)_2_:AK2 complex simulations, and dimeric AIFM1(NAD+FAD):AK2, taken from the AIFM1(NAD+FAD)_2_:AK2 complex simulations, respectively

The x-axis represents the interacting pairs (where chain A corresponds to AIFM1 and chain B corresponds to AK2), while the y-axis indicates the persistence rates of these salt bridges. Light-coloured sections of the stacked bars denote the percentage of salt bridge interactions formed during one 2 µs long MD simulation, whereas dark-coloured sections represent the corresponding replica 2 µs long MD simulation. For each AIFM1 scenario, C-loop interactions persisting more than 20% of the simulation time were considered. Except for a few consistent pairs, most salt bridge interactions within the AIFM1 C-loop exhibit significant variability between replicated MD simulations. This suggests that these salt bridges are non-specific and can be formed by multiple possible interacting pairs. This instability likely contributes to a destabilized conformation and the dynamic nature of the C-loop. The number of interacting pairs for each case correlates well with the extent of the resolved regions of AIFM1 (see Fig. 3).

**Supplemental Movie. The representative dynamic behaviour of the AIFM1(NAD+FAD)_2_:AK2 complex as observed from 0.5 to 1.4 µs in one replica MD simulation.**

The movie demonstrates the swinging motion of AK2 when bound to AIFM1(NAD+FAD), as captured by our MD simulations. In the movie, AK2 is depicted in blue, the AIFM1 dimer is in grey, and the C-loops are highlighted in orange. Towards the end of the movie, the negatively charged residues between 533 and 540 (ETESEASE) of the AIFM1 C-loop start interacting with the positively charged AK2 helix residues between 62 and 72 (KKLKATMDAGK). This interaction showcases how the electrostatic attraction between these charged regions facilitates the AIFM1/AK2 interaction.

### Mutational analysis of AK2 recruitment

The binding mechanism formulated here requires C-loop release but not dimerisation of AIFM1. To test this, a triple mutant in the dimerisation interface was prepared, rendering AIFM1 obligate monomeric but still permissive to NADH-induced C-loop release^49,50^. This variant was unable to form dimers in the presence of NADH, with only a slight peak shift towards higher apparent molecular weight, likely attributable to C-loop order-to-disorder transition (Fig. 3A, Fig. 4A, blue and green traces). However, AK2 was recruited in a NADH-dependent manner (Fig. 4A, grey and orange traces), with mass photometry suggesting a 1:1 stoichiometry (Fig. S6). Another AIFM1 variant, Cβ-clasp W196A mutant, shows constitutive C-loop detachment and dimerisation, independent of NADH-loading^49^. Accordingly, the SEC elution volume was shifted towards dimeric AIFM1 in the absence of NADH with only a slight further shift upon NADH addition (Fig. 4B, blue and green traces). As predicted, this AIFM1 variant was able to bind AK2 regardless of NADH presence (Fig. 4B, grey and orange traces, Fig. S6). Thus, C-loop release and not AIFM1 dimerisation is the critical determinant for AK2 recruitment.

**Fig. 4:**
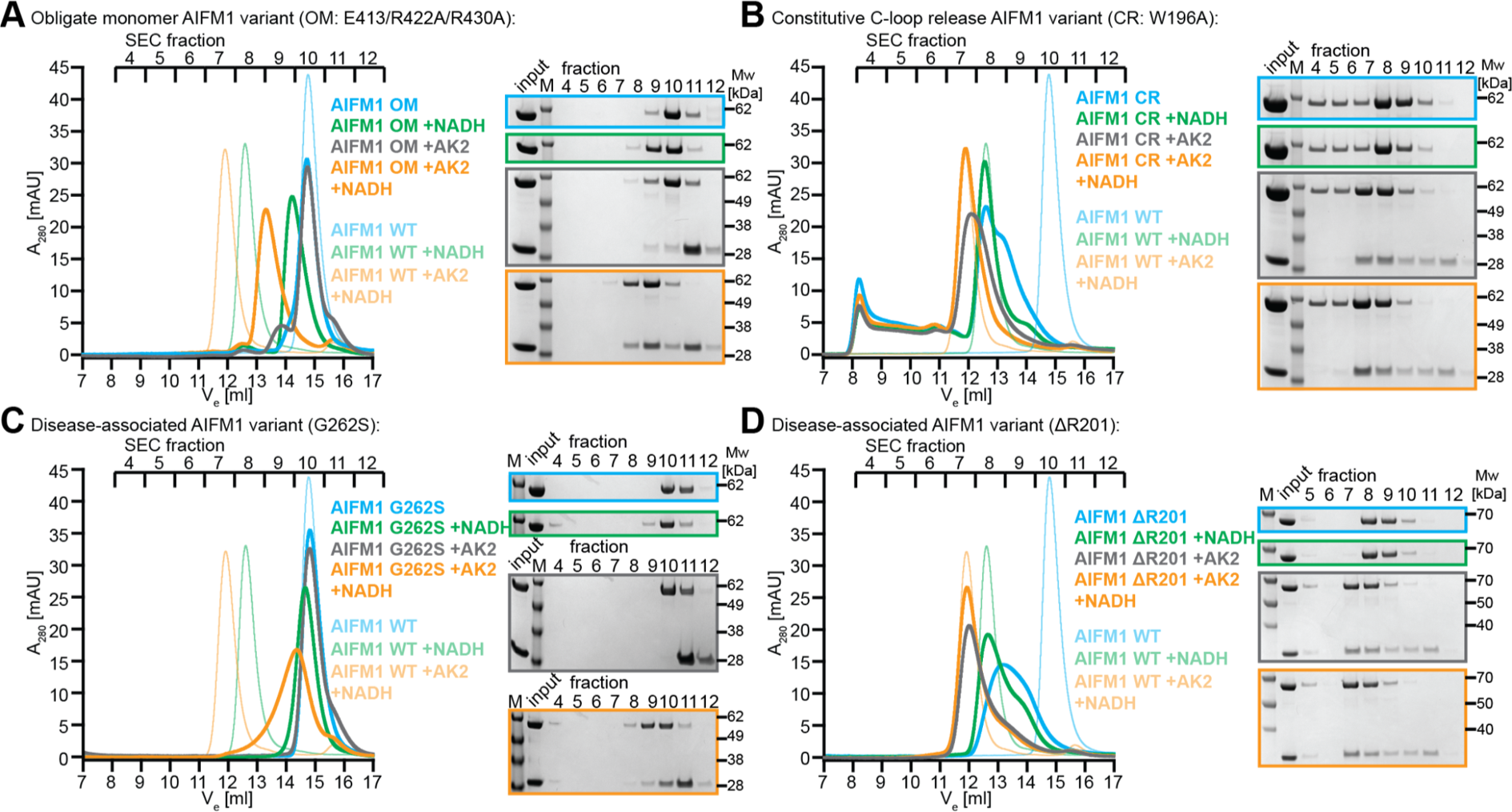
Structure-based and disease-associate mutations impinge on the AIFM1/AK2 interaction. (A) *In vitro* reconstitution, followed by SEC, of an obligate monomeric AIFM1 variant (OM), AK2, and an equimolar mixture of both proteins, in the absence and presence of NADH. WT traces from Fig. 2A are shown in pale colours for comparison. (B) *In vitro* reconstitution, followed by SEC, of a constitutive C-loop release AIFM1 variant (CR) as in A. (C) *In vitro* reconstitution, followed by SEC, of a disease-related AIFM1 variant (G262S) as in A. (D) *In vitro* reconstitution, followed by SEC, of a disease-related AIFM1 variant (ΔR201) as in A.

**Fig. S6:**
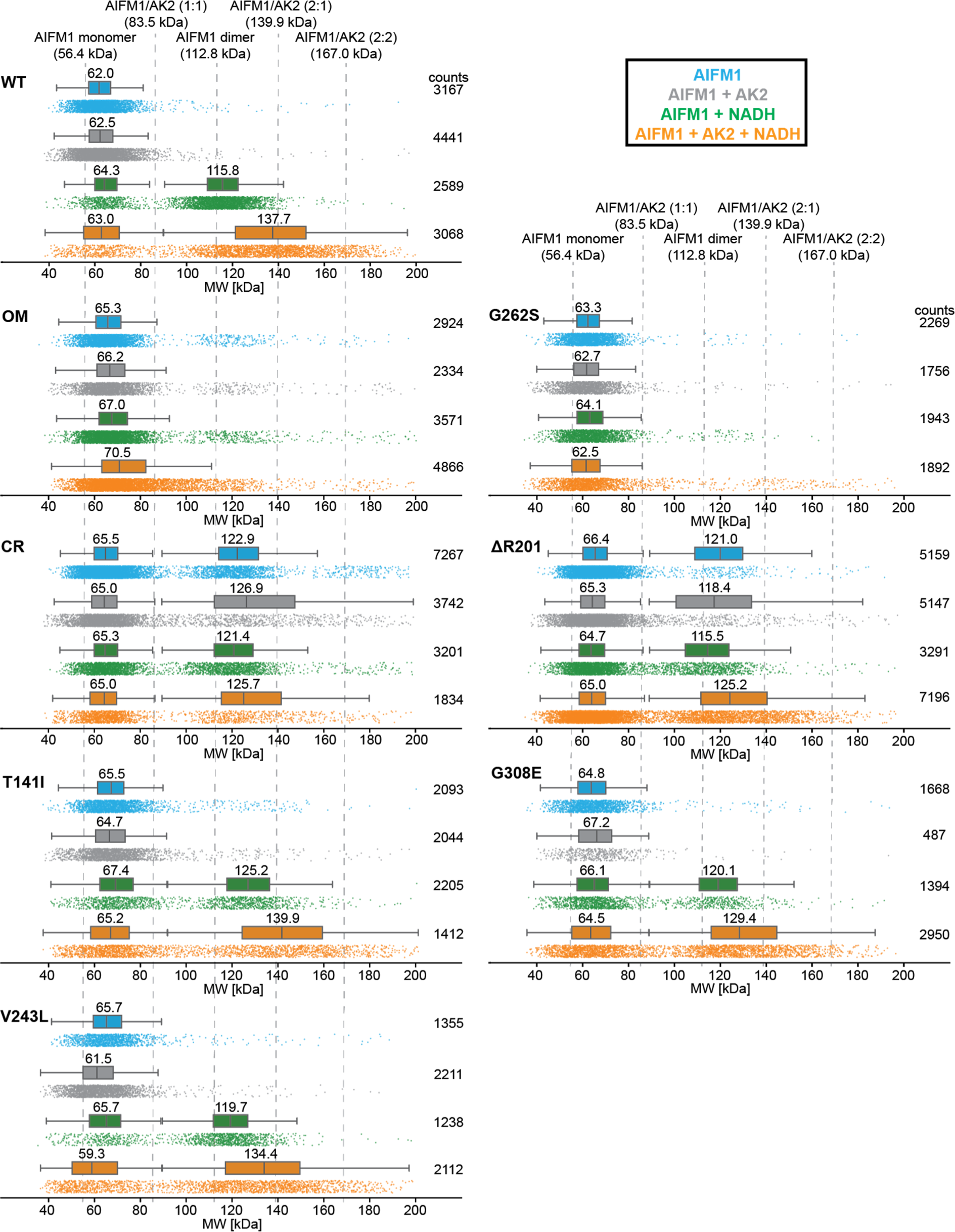
Additional mass photometry analyses of AIFM1 variants in the absence and presence of NADH and AK2, related to Fig. 4. Box-and-whisker plots of apparent molecular weights are shown above the raw data counts, with the median molecular weights indicated. Box-and-whisker plots are drawn for molecular weight ranges between 1 kDa and 90 kDa, and if applicable between 90 and 200 kDa, except for AIFM1 OM, where the range was 1 kDa to 140 kDa. AIFM1 WT data from Fig. 2B are shown on top for comparison. Theoretical molecular weights of AIFM1 and AIFM1/AK2 assemblies are shown on top of the uppermost graph and indicated by the grey dashed lines.

### Disease-causing AIFM1 mutations and AK2 recruitment

We next investigated disease-causing AIFM1 mutants that disrupt the allosteric pathways of AIFM1 at different points. The first variant, AIFM1 G262S, was identified in a patient suffering from slowly progressive mitochondrial encephalomyopathy. G262S expression levels were reduced by half in patient fibroblasts, and muscle homogenates showed an 80% reduction in OXPHOS CIV activity^53^. Previous *in vitro* characterisation of recombinant G262S revealed perturbations of the NADH-binding site and highly reduced NADH affinity^54^. Consistently, the G262S variant in the presence of NADH remained monomeric (Fig. 4C, blue and green traces, Fig. S6) and was impaired in AK2 binding (Fig. 4C, grey and orange traces, Fig. S6). This is in contrast to the obligate monomeric but AK2-binding competent AIFM1 OM variant (Fig. 4A, orange trace, Fig. S6). These data indicate that cofactor-binding site disturbances in G262S block allosteric pathways to both, dimer formation and C-loop release, the latter preventing AK2 recruitment.

The second disease-associated variant tested was AIFM1 ΔR201, found in two infant patients suffering from fast-progressing, severe mitochondrial encephalomyopathy and OXPHOS failure^55^. ΔR201 performed very similarly to the constitutive C-loop release AIFM1 variant W196A, dimerising even in the absence of NADH (Fig. 4D, blue and green traces, Fig.S6), and constitutively recruiting AK2 (Fig. 4D, grey and orange traces, Fig. S6). Given the location of R201 in AIFM1’s Cβ-clasp region, close to W196, it is tempting to speculate that deletion of R201, concomitant with loss of the electrostatic contact to the C-loop residue E531 (Fig. 3A), triggers constitutive C-loop release, rationalising the W196A phenocopy, i.e. NADH-independent AIFM1/AK2 interaction. Taken together, we demonstrate that the two disease-associated AIFM1 mutations examined above have strong and opposing effects on AIFM1/AK2 interaction *in vitro*. However, additional disease-related AIFM1 mutations that we tested showed no overt phenotype on AK2 recruitment (Fig. S6), implying pleiotropic AIFM1-dependent pathogenesis mechanisms.

### Glycolysis inhibition curtails AIFM1/AK2 association

Next, we aimed to test our model of NADH-dependent AIFM1/AK2 interaction in a cell culture setting. Cellular NADH is generated primarily by two processes: the TCA cycle in the mitochondrial matrix and glycolysis in the cytoplasm. The presence of solute-permeable porins in the mitochondrial OM allows for the exchange of metabolites, while the IM is tightly sealed^56,57^. As a result, NADH levels in the mitochondrial IMS, where the NADH-binding domain of AIFM1 is located, are largely influenced by glycolytic NADH production and active transport out of the mitochondrial matrix (MM) through shuttle mechanisms^58^.

To investigate the impact of inhibiting processes that maintain IMS NADH availability on AIFM1/AK2 interaction, we inhibited glycolytic NADH production using the GAPDH inhibitor iodoacetate (IA^59^) in 293T cell cultures. Indeed, this intervention reduced the affinity of AIFM1 to AK2, as confirmed by affinity purification of ectopically expressed FLAG-AK2, followed by western blotting (Fig. 5A) and quantitative AP-MS (Fig. 5B). Thus, glycolytic NADH production is a driver of AK2 recruitment by AIFM1, in line with our mechanistic observations.

**Fig. 5:**
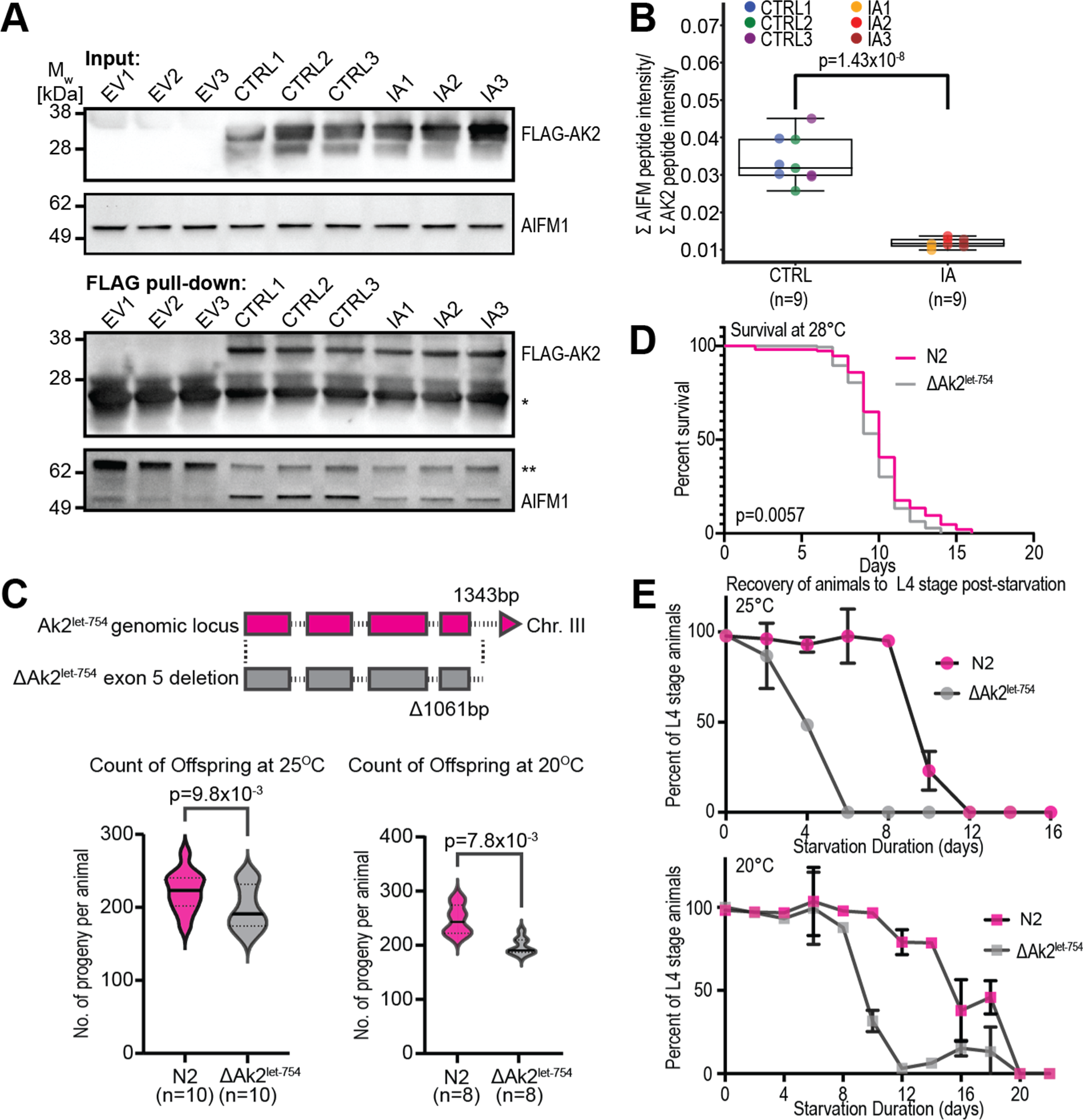
Cellular and organismal analyses of the AIFM1/AK2 interaction. (A) Immunoblot analysis of cell lysates and FLAG pull-downs of 293T cells, transfected with empty vector (EV) or FLAG-AK2, either untreated (CTRL) or treated with the GAPDH inhibitor iodoacetate (IA). *, ** - non-specific labelling by the secondary antibody. (B) MS-based label-free quantification of the ratio of AIFM1/AK2 peptide intensities from CTRL or IA-treated FLAG pull-downs. (C) (Upper panel) Schematic of the genetic targeting strategy to delete exon 5 (amino acid residues 246-251) from the endogenous *C. elegans* AK2 (*let-754*) locus. (Lower panel) Brood size assay comparing WT (N2) and AK2 exon 5 deletion (ΔAk2^let-754^) animals at the indicated temperatures. (D) Lifespan assay at 28°C, comparing WT (N2) and AK2 exon 5 deletion (ΔAk2^let-754^) *C. elegans animals.* (E) L1 recovery assays comparing WT (N2) and AK2 exon 5 deletion (ΔAk2^let-754^) *C. elegans* animals at the indicated temperatures.

### C-terminal truncation of AK2 reduces organismal fitness

To test the physiological relevance of NADH-dependent AIFM1/AK2 interaction, we turned to *C. elegans* as a model organism. Importantly, the AIFM1-interacting C-terminal motif is conserved in *C. elegans* AK2 (Fig. S7A), and AlphaFold3 structure prediction suggested a similar β-strand insertion mechanism in this species (Fig. S7B). Accordingly, we generated a *C. elegans* mutant where the sole AK2 homologue *let-754* lacks the C-terminal six amino acid residues representing the AIFM1 binding motif (ERVSFV 246-251) by removing the final exon from its coding sequence (Fig. 5C, Fig. S7A). These animals were of superficially wild-type appearance. We therefore designed three experiments based on our *in vitro* and cell-based understanding of the AK2 C-terminus’ role in directing ATP usage.

**Fig. S7:**
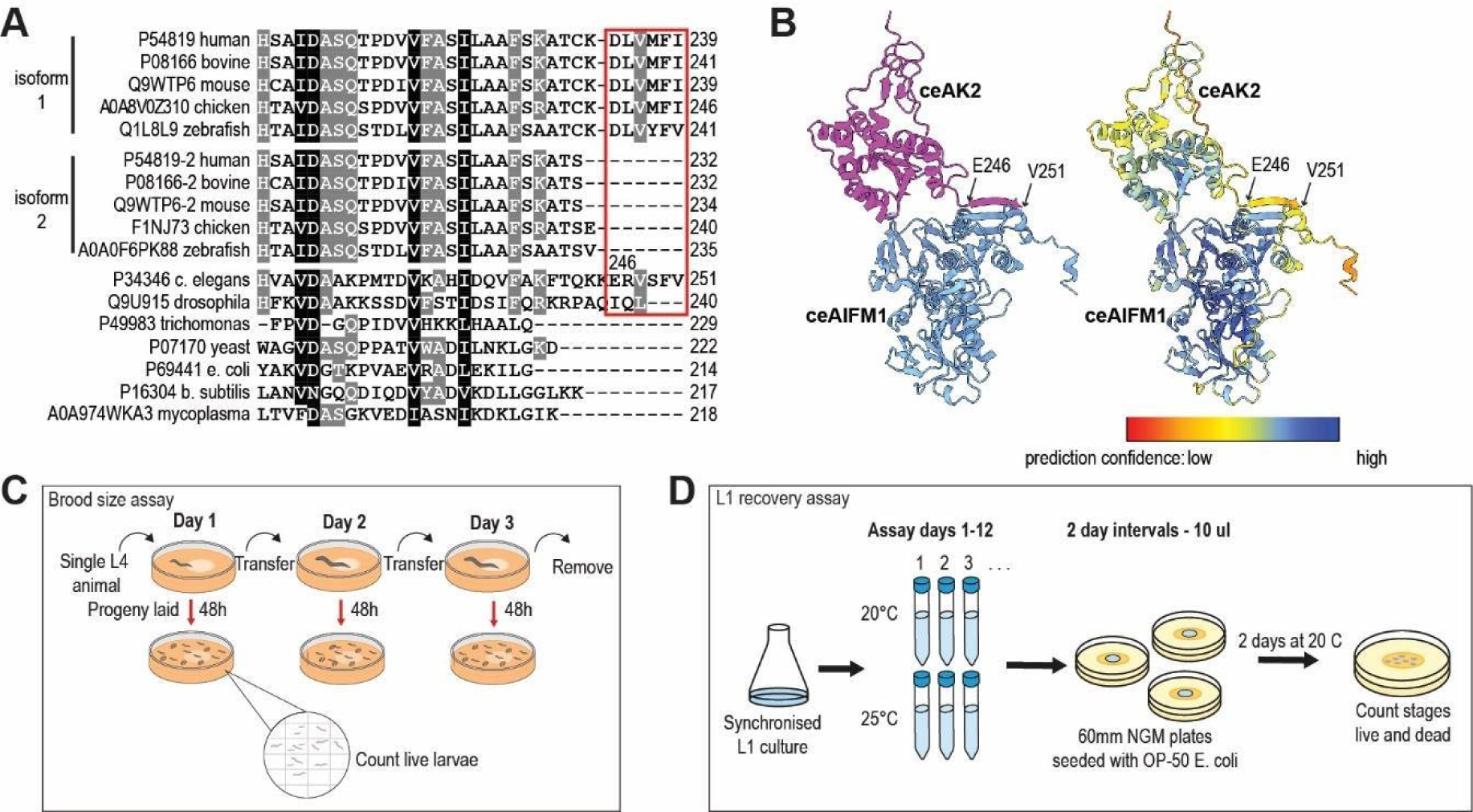
*C. elegans* experiments, related to Fig. 5. (A) Multiple sequence alignment of AK2 homologues. Where applicable, isoforms 1 and 2 are grouped separately. UniProt accession numbers are given on the left. The red box indicates the amino acid stretch corresponding to the C-terminal AIFM1-interacting β-strand motif. (B) The amino acid sequences of *C. elegans* AIFM1 (UniProt ID Q9U229, 2 copies) and AK2 (UniProt ID P34346, 1 copy) were used for AlphaFold v3 structure prediction together with 2 copies of NADH and FAD, respectively^66^. The resulting model is shown in cartoon representation, in the left panel coloured according to protein identity (AIFM1 - blue, AK2 - pink) and in the right panel according to prediction confidence in a colour gradient from lowest confidence (pLDDT score 0) in red to highest confidence (pLDDT score 100) in dark blue. The second AIMF1 protomer is hidden for clarity. (C) Schematic of the *C. elegans* brood size assay workflow. (D) Schematic of the *C. elegans* L1 recovery assay workflow.

As fertility is tightly associated with mitochondrial energy metabolism^60–62^, and interestingly also affected in AIFM1 knockout *C. elegans* animals^63,64^, we counted the number of viable offspring produced by these mutants and found that they laid significantly fewer eggs developing into adults than wild-type control animals (Fig. 5C, Fig. S7C). Since the *C. elegans* life cycle is highly temperature dependent, we counted the brood size at both 20°C and 25°C: at both temperature conditions, fewer viable eggs were laid by the mutant. These data show that an intact AK2 C-terminus, which is required for recruitment to AIFM1, is necessary to sustain normal fertility in *C. elegans*.

The lifespan of *C. elegans* inversely correlates with the temperature the animals are kept at, because their metabolic rate rises with temperature^65^. Increased metabolic rates, concomitant with higher glycolytic activity, should stimulate AIFM1/AK2 interaction. Consequently, loss of the AK2 C-terminus should compromise the mutant animals upon cultivation at elevated temperatures. Indeed, in life span assays at 28°C, we found the mutants’ lives to be significantly shorter than that of wild-type animals (p = 0.0057, Fig. 5D).

For similar reasons, the ability of the mutant animals to cope with fluctuating nutrient availability should be adversely affected. Wild-type and mutant animals were arrested in larval stage 1 (L1) by starvation for varying periods, then transferred back to a normal growth medium, and after 48 h the percentage of animals that progressed to larval stage 4 (L4) was scored (Fig. S7D). Again, the experiments were conducted at 20°C and 25°C temperature. Under both conditions, a marked difference in the recovery from L1 arrest between wild-type and mutant animals was observed, increasing in magnitude in dependence on the length of the starvation period (Fig. 5E). Accordingly, an intact AK2 C-terminus seems also critical for starvation stress resistance and the transition from starvation-induced L1 arrest (low glycolytic activity) to growth under nutrient-rich conditions (high glycolytic activity).

## Discussion

AK2 has been indirectly linked to AIFM1 as a substrate of the mitochondrial disulfide relay import pathway, in which AIFM1 plays a central role by providing a binding platform for the oxidoreductase MIA40/CHCHD4^30–34^. Here, we provide evidence for stable, NADH-dependent recruitment of AK2 by AIFM1 also in the absence of MIA40. In addition, direct AIFM1 association with OXPHOS CIV has been described previously but the functional implications remained undefined^67,68^. Our data suggest a model where AIFM1 acts as a cellular NADH sensor, to position AK2, under conditions of high NADH availability, close to OXPHOS/ATP synthase protein complexes and ANTs. There, AK2 is properly placed to locally regenerate ADP as a substrate for ATP synthesis, using ATP that emanates from ANTs and generating fresh ADP for return to ATP synthase. This system supports ATP production by a stable supply of ADP substrate to fully utilise OXPHOS capacity (Fig. 6, left panel). This provides a feed-forward mechanism during transitions to increased glycolytic flux and a cellular adaptation to elevated ATP synthesis needs that commit a proportion of fresh ATP to fill the ADP pool. Under starvation conditions, where glycolysis and consequently NADH levels decrease, AK2 is decoupled from AIFM1, and accordingly, local ATP consumption/ADP generation by AK2 is suspended (Fig. 6, right panel). In conclusion, our model represents a metabolic, NADH-driven switch distributing ATP flow between investment into ADP build-up and cellular consumption.

**Fig. 6:**
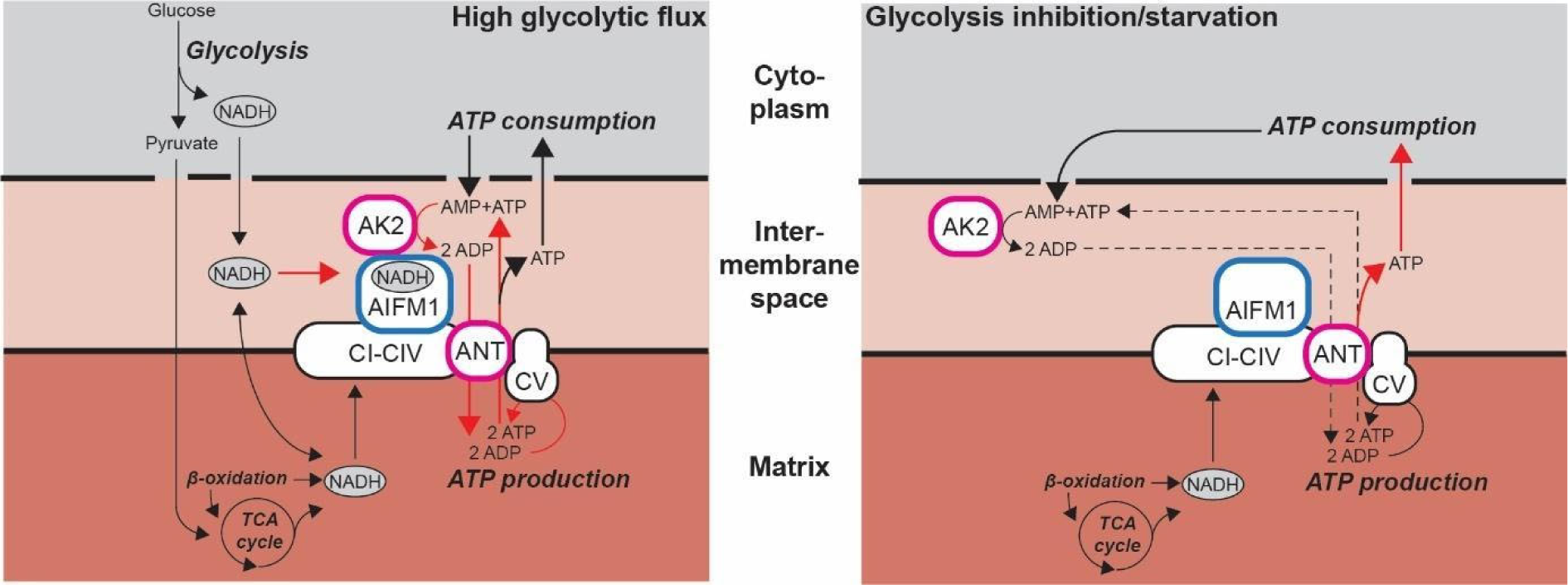
Schematic of the AIFM1/AK2 module’s proposed gatekeeping role in cellular energy metabolism. (A) (Left panel) High glycolytic flux leads to elevated cellular NADH levels, boosting the electrochemical gradient and ATP synthesis. In anticipation of the increased need for ADP as a substrate of ATP synthesis, AIFM1 senses NADH and recruits AK2 to the vicinity of OXPHOS complexes and ANT to direct ATP flux into regenerating ADP. (Right panel) When glycolysis is inhibited or during starvation, cytoplasmic and IMS NADH levels decrease, weakening the AIFM1/AK2 interaction and de-localising AK2. This results in longer diffusion pathways and slower ATP use by AK2, conserving freshly synthesised ATP for cellular use. In essence, when NADH levels are high, ADP regeneration is prioritised; when NADH levels are low, ADP regeneration is deprioritised.

An astonishingly high number of mutations in AIFM1 are linked to inherited mitochondrial diseases^37,69–73^, and the most common causes of mitochondrial diseases are OXPHOS defects^11,74^. In line with AIFM1’s role in the mitochondrial disulfide relay, disease-associated mutations causing AIFM1 knockout or hypomorphism interfere with the import and oxidative folding of disulfide relay import substrates including ETC components, leading to OXPHOS dysfunction^33,34,36,75^. Our biochemical findings imply two AIFM1 disease-related mutations in either inhibition or constitutive activation of AK2 recruitment (Fig. 4C, D). According to our model, both opposing activities might converge on disturbances of CV substrate supply, impinging on cellular energy metabolism, thus offering an additional pathogenesis mechanism of inherited AIFM1 mutations.

The C-terminal extension of AK2 that binds to AIFM1 coincides evolutionary with the arrival of multicellularity, being observed in vertebrates and nematodes like *C. elegans* but not bacteria and yeast (Fig. S7A). In vertebrates including humans, two splice variants of AK2 have been described^26,76^. They differ in sequence only in the C-terminus, where in isoform 2 (iso2) the last eight amino acids are replaced by a single serine residue. Interestingly, the amino acid stretch deleted in iso2 exactly corresponds to the region we have identified as crucial for NADH-dependent AIFM1 interaction (Fig. 2D-F). This presents an opportunity for the regulation of AIFM1/AK2 interaction by alternative splicing. Indeed, in humans, there are tissue-specific differences in the ratio of AK2 iso1/iso2 expression observable, with the highest values in muscle and male reproductive tissues, in line with fluctuating energy demands in these tissues^77–80^, and low values in brain tissue (Fig. S8). This supports a possible link between AK2 recruitment and the clinical presentation of AIFM1 mutants as encephalomyopathies.

**Fig. S8:**
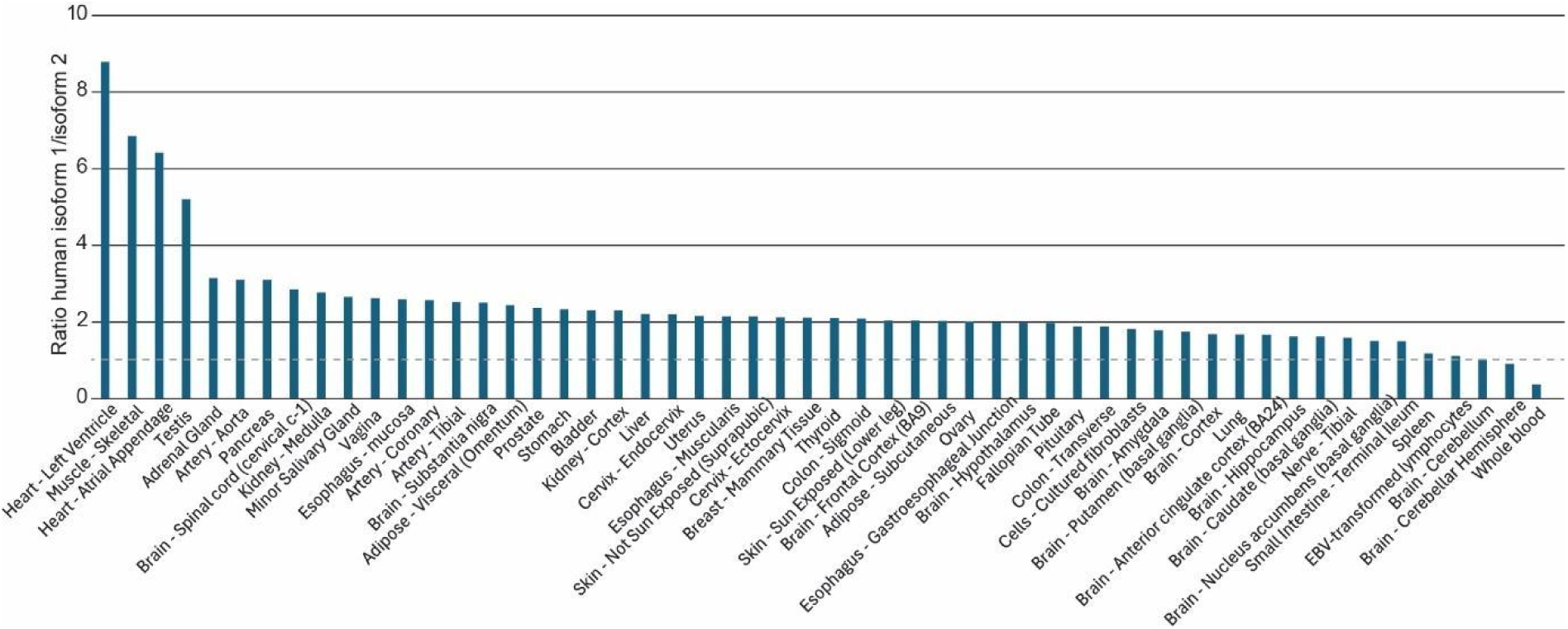
Tissue-specific expression of human AK2 isoforms, related to Fig. 6. The tissue-specific ratio of AK2 isoform 1/isoform 2 expression has been calculated by division of the corresponding transcript-per-million values obtained from the Genotype-Tissue Expression (GTEx) Portal Transcript Browser.

In conclusion, we here combined *in situ* crosslinking-based PPI network analysis with biochemical, structural, computational, cellular biology, and genetics to pinpoint a mechanistic link between NADH sensing, ATP synthase substrate supply and ATP flux regulation in multicellular species. In the broader context, our work exemplifies the concept of substrate supply logistics as a general biological organisation principle to ensure efficient and controlled catalysis. Communication between cellular compartments requires tightly coordinated and consequently, controllable machinery, akin to “just in time” logistics, guaranteeing favourable substrate concentrations and product consumption, attuned to cellular needs.

## Methods

### Plasmids

The expression plasmids encoding human AIFM1 (residues 102-613), AIFM1 102-613 variants, and AK2 were codon-optimised for *E. coli*, synthesised and ligated in the pET151/D-TOPO backbone by GeneArt (Thermo Fisher Scientific). The expression plasmid encoding full-length human AIFM1 was synthesised and ligated in the pcDNA3.1(+) backbone by GeneArt (Thermo Fisher Scientific). Insert sequences are provided in the Supplemental Material.

### Cell lines

*E. coli* BL21(DE3) were obtained from New England Biolabs (cat# C2527H) and cultivated in LB medium under shaking at 37°C or 18°C as described below (protein expression and purification).

K-562 cells (DSMZ, cat# ACC-10) were grown at 37°C under a humidified atmosphere holding 5% CO2 in RPMI 1640 medium (Corning) containing 10% fetal bovine serum (Sigma-Aldrich cat# S0615-500ML) and 2 mM L-glutamine (Corning cat# 25-005-CI).

293T cells (DSMZ, cat# ACC-635) were grown at 37°C under a humidified atmosphere holding 5% CO2 in DMEM (Corning, cat# 15-013-CV) supplemented with 10% FBS (Sigma-Aldrich, cat# S0615-500ML) and 2 mM L-glutamine (Corning, cat# 25-005-CI).

Both, K-562 and 293T cells tested negative for mycoplasma contamination.

### Mitochondria isolation and crosslinking

Per mitochondria preparation, 4 x 8.75 million K-562 cells were seeded 3 days in advance in T175 suspension culture flasks (650 mL, cellstar®, Greiner Bio-One). On the day of mitochondria preparation, ca. 200-300 million K-562 cells were collected by centrifugation (5 min at 200 x g, Mega Star 600 R centrifuge (VWR), rotor #75005701 (Thermo Scientific)) and washed twice with phosphate-buffered saline (Dulbecco’s PBS, without Mg2+/Ca2+, Corning). Cell lysis and subsequent mitochondria preparation were performed on ice. Briefly, cell lysis was carried out in 5.5 mL ice-cold RSB hypotonic buffer (10 mM NaCl (Sigma), 1.5 mM MgCl2 (Sigma), 10 mM HEPES (Sigma), pH 7.5) using Dounce homogenization. By 5-8 strokes of the piston in a 7 mL Dounce tissue grinder (Wheaton) cells were lysed until approx. 20% of the cells were still intact to avoid nuclei damage. 4 mL ice-cold 2.5 x MS homogenisation buffer (525 mM mannitol (Roth), 175 mM sucrose (Sigma), 2.5 mM EDTA (Sigma), 12.5 mM HEPES, pH 7.5) was added to obtain an isotonic solution. To clarify the cell lysate, it was centrifuged 3-5x at 1,300 x g (5 min, 4 °C, Heraeus multifuge X3R, Thermo Scientific, rotor F15-8×50cy) until no pellet was visible. The mitochondria were pelleted by centrifugation at 12,400 x g (15 min, 4 °C, centrifuge as before) and washed once with 5 mL ice-cold 1x MS homogenization buffer. The isolated mitochondria were resuspended in 20 mM sodium phosphate pH 8.0, 150 mM NaCl (PBS, Sigma). The protein concentration was estimated via Bradford Assay according to the manufacturer’s protocol (BioRad, #500-0006).

Disuccinimidyl sulfoxide (DSSO) and anhydrous dimethylformamide (DMF) were purchased from Thermo Scientific Pierce. Isolated mitochondria were washed in ice-cold PBS and pelted at 12,400 x g (15 min, 4 °C, as above). 5 mg proteins were chemically crosslinked using 0.48 mM DSSO in DMF which equals a protein-to-crosslinker ratio of 6:1 at 1 mg/mL protein concentration in 5x 1.0 mL crosslinking reactions. After 1 h incubation on ice, the crosslinking reaction was quenched by adding ammonium bicarbonate to a final concentration of 50 mM (30 min on ice). After crosslinking, a further centrifugation step (12,400 x g, 15 min, 4 °C) was performed to pellet intact crosslinked mitochondria. The resulting pellet was dissolved in 2 mL PBS and mitochondria lysis was achieved by sonication (6×10 sec., duty cycle 50%, output 1, on ice, Sonifier 250 (Branson) equipped with Ultrasonicator microtip (Branson)).

### Proteolytic digestion

After DSSO crosslinking, centrifugation at 20.000 x g (30 min, 4 °C, Eppendorf centrifuge #5804R, rotor FA-45-30-11) separated the proteins into a soluble supernatant and an insoluble pellet. The proteins in the separated supernatant and the pellet proteins were precipitated by adding the 4 x volume of ice-cold acetone (Sigma), harsh vortexing (and 10 min sonication bath to resolve the pellet) and incubating overnight at −20°C. Proteins were pelleted by centrifugation at 15.000 x g (10 min, 4°C in cold cabinet, Eppendorf centrifuge #5418, rotor FA-45-18-11). After allowing the pellet to air dry, samples were tryptic digested. The crosslinked proteins were denatured using 6 M urea (Sigma), 2 M thiourea (Sigma) in 100 mM ammonium bicarbonate. The insoluble pellet was digested with the help of 0.1% (v/v) RapiGest SF Surfactant (Waters) in 100 mM ABC. Disulfide bonds were reduced with 10 mM dithiothreitol (DTT, Sigma) for 30 min at 25 °C, gentle agitation. To alkylate reduced disulfide bonds, 40 mM iodoacetamide (IAA, Sigma) was added and incubated for 30 min at 25 °C, with gentle agitation in the dark. By adding 10 mM DTT the alkylation reaction was quenched. Note that the DTT/IAA concentration was estimated in dependency of the protein concentration (5 mM DTT for reduction and quenching reaction per 1 mg/mL protein and 15 mM IAA per 1 mg/mL protein).

The pre-digestion was performed in 200 μg portions by adding 1:100 LysC (Thermo Scientific) and incubating 4 h at 25 °C, gentle agitation. After diluting with 100 mM ammonium bicarbonate to a final concentration of 1.5 M urea/thiourea, trypsin (Thermo Scientific) was added at an enzyme-to-substrate ratio of 1:50 and incubated for at least 16 h (25 °C, gentle agitation). The in-solution-digestion was stopped by adding 10% (v/v) trifluoroacetic acid (TFA, Thermo Scientific) till reaching pH ≤ 2. To get rid of RapiGest SF Surfactant, the insoluble sample of DSSO-crosslinked peptides was incubated 1 h at 25 °C and clarified by spinning down twice at 14.500 x g, 10 min (Eppendorf centrifuge #5418, rotor FA-45-18-11).

### Peptide Purification

Peptide desalting and purification were performed after tryptic digestion of DSSO-crosslinked proteins using EmporeTM Solid Phase Extraction Cartridges C18-SD according to the manufacturer’s protocol. Centrifugation steps were carried out for 1 min (45 x g, room temperature, Eppendorf centrifuge #5804R, swing out rotor A-4-44). Briefly, the C18 membrane was activated by adding 1 mL methanol (MeOH), spin and washed with 500 μL 80% (v/v) acetonitrile (ACN, Sigma) in 0.1% (v/v) TFA. The acidified tryptic peptides were loaded with a maximal load of 750 μg per cartridge. After washing with 0.1% (v/v) TFA, elution was accomplished by adding 150 μL MeOH and 175 μL 80% (v/v) ACN in 0.1% (v/v) TFA. Then, the samples were divided into portions of 400 μg tryptic peptides each, evaporated completely in an Eppendorf vacuum concentrator and stored at −20 °C until chromatographic fractionation. Peptide desalting and purification were performed after SCX and hSAX fractionation of DSSO-crosslinked proteins using EmporeTM C18 Extraction Discs (3M). This StageTip purification was based on the previously published protocol^81^ and all steps were carried out using GilsonTM GMCLabTM Fixed-Speed Mini Centrifuge (Thermo Scientific). Briefly, the C18 membrane was activated using 20 μL LC-MS grade MeOH (VWR), cleaned by 20 μL 80% (v/v) ACN in 0.1% (v/v) TFA and equilibrated using 2×20 μL 0.1% (v/v) TFA. After sample loading with maximal 15 μg peptides, the StageTips were washed twice using 20 μL 0.1% (v/v) TFA. Purified peptides were stored on StageTips either at −20°C or eluted directly using 80% (v/v) ACN in 0.1% (v/v) TFA.

### Strong Cation Exchange (SCX) Chromatography

DSSO-crosslinked peptides were fractionated using strong cation exchange chromatography (SCX) as previously described in Chen et al., 2010. In our workflow, aliquots of 400 μg peptides were resuspended in 130 μL SCX buffer A (30% (v/v) ACN, 10 mM KH2PO4, pH 3.0,). 120 μL peptide sample was loaded onto a PolyLC Polysulfoethyl ATM column (100 x 2.1 mm, 3 μm, 300 Å) operated at room temperature under Unicorn 7.1 on an Äkta pure HPLC system. Peptide separation was achieved using a non-linear gradient with SCX buffer B (30% (v/v) ACN, 1 M KCl, 10 mM KH2PO4, pH 3.0) at a constant flow rate of 0.15 mL/min. Fractions of 200 μL each were collected over the elution window resulting in 11 SCX fractions of interest in total. Collected SCX fractions from four to five runs were pooled SCX fraction-wise, desalted using StageTip purification like previously published^81^ with four C18 StageTip discs per tip and a maximum load of 15 μg and stored at −20°C.

### Hydrophilic Strong Anion Exchange (hSAX) Chromatography

The desalted peptides were fractionated by hydrophilic strong anion exchange (hSAX) chromatography using a resin-based Dionex IonPac AS24 column (2 x 250 mm) operated at 15°C under Unicorn 7.1 on Äkta pure HPLC system. StageTip cleaned and evaporated samples were resuspended in 220 μL hSAX buffer A (20 mM Tris-HCl, pH 8.0 at 15 °C), injected and separated at a constant flow of 150 μL/min as previously published^82^. Elution of peptides was accomplished by an exponential gradient with increasing salt concentration in hSAX buffer B (1 M NaCl, 20 mM Tris-HCl, pH 8.0 at 15 °C). 150 μL fractions were collected and pooled to 10 hSAX samples per SCX fraction. Peptides were acidified with TFA to pH 2, desalted using StageTip purification as described above and stored at −20°C.

### Crosslinking mass spectrometry (CLMS)

In total, 110 different SCX-hSAX fractions per insoluble and soluble proteome were analysed using an UltiMate 3000 Nano LC system coupled to a Q Exactive HF Hybrid Quadrupole-Orbitrap mass spectrometer (Thermo Fisher Scientific, San Jose, USA). Completely evaporated samples were resuspended in 1.6% (v/v) ACN and 0.1% (v/v) formic acid (FA) and iRT peptides (Biognosys) were added 1:20 according to manufacturing instructions. Generally, SCX-hSAX fractions with >24 mAU in hSAX chromatography dimension were injected as technical duplicates, fractions with higher sample amounts even as triplicates. A minority of fractions with lower sample amounts were single-injected or even pooled (primarily hSAX fractions 3-4 and 9-10). Consequently, 246 MS runs were acquired for insoluble and 177 for soluble mitochondrial samples. The reverse phase chromatography was running at a constant flow rate of 250 nL/min with mobile phase A containing 0.1% (v/v) FA in water and mobile phase B 80% (v/v) ACN and 0.1% (v/v) FA in water. Peptides were injected onto a 500 mm C-18 EasySpray column (75 μm ID, 2 μm particles, 100 Å pore size) and eluted usually in a gradient from 2% – 7.5% – 42.5% – 50% mobile phase B for 87 min^82^. Note that very hydrophobic peptides or very hydrophilic peptides (in early/late SCX-hSAX fractions) were run by adapted LC-gradient to higher and/or lower phase B concentrations.

Here, MS1 spectra were obtained at 120,000 resolution (orbitrap, AGC target 3 × 106 ions and maximum injection time of 50 ms). Precursor ions with charge states 3-6 and an intensity higher than 4.2 × 103 were isolated for stepped CID-based fragmentation (using collision energy of 18, 24 and 30, isolation window 1.4 m/z, AGC target 5 × 104, 120 ms maximum injection time). MS2 spectra were recorded at 60,000 resolution in the orbitrap. The non-fractionated aliquots for protein identification in the soluble and insoluble proteome were run as triplicate over an adapted 180-minute gradient and without discriminating lower charge states for MS2 selection.

### Crosslinking database search and FDR

Raw mass spectrometry data were processed using msConvert (version 3.0.11729) to convert the data to MGF format. Precursor and fragment masses were recalibrated to account for any mass shifts during measurement. The resulting peak files were analyzed using xiSEARCH 1.7.6.4 with the following parameters: MS1/MS2 error tolerances set at 2 and 5 ppm respectively, allowing up to 2 missing isotope peaks. Tryptic digestion specificity was used with up to 2 missed cleavages, and carbamidomethylation on cysteine was set as a fixed modification. Variable modifications included oxidation on methionine, methylation on aspartic and glutamic acid, deamidation on asparagine, pyroglutamate on glutamine, and crosslinker modifications (“DSSO-NH2” 175.03031 Da, “DSSO-OH” 176.01433 Da) on lysines. Losses considered were –CH3SOH, –H2O, and –NH3. Additional loss masses for crosslinker-containing ions were defined to account for the cleavability of DSSO (“A” 54.01056 Da, “S” 103.99320 Da, “T” 85.98264 Da). Matches were filtered to include at least one peptide with a DSSO doublet. Crosslink sites were allowed on the side chains of Lys, Tyr, Ser, Thr, and the protein N-terminus. A non-covalent crosslinker with a mass of zero was used to flag spectra potentially arising from gas-phase-associated peptides, which were removed before false-discovery-rate (FDR) estimation. Crosslinks containing a variable modification were placed in a separate group during FDR calculation. The database was based on linear acquisition from 1e6, with transit peptides removed. Results from soluble and insoluble searches were normalized by CSM-level FDR, and then FDR was calculated as described for *E. coli*^82^, with a threshold of 5% for link level and 2% for PPI level.

### Blue native polyacrylamide gel electrophoresis (BN-PAGE)

Mitochondria were isolated from K562 cells as described above and flash-frozen in 1x MS homogenization buffer (210 mM mannitol, 70 mM sucrose, 5 mM HEPES pH 7.5, 1 mM EDTA). After thawing, mitochondria were pelleted at 15,000 x g for 15 min at 4°C and resuspended in PBS at 400 µg/mL. DSSO crosslinking of mitochondria was performed at 0.25 mM final concentration (using a 50 mM freshly prepared stock in anhydrous DMF) for 30 minutes at 25°C while gently rotating. A solvent-control (0 mM DSSO, Fig. S2C, left) was performed in parallel. The crosslinking reaction was quenched by incubation with Tris pH 8.0 (20 mM final concentration) for 30 minutes at 25°C while gently rotating.

80 µg DSSO-crosslinked mitochondria were pelleted at 15,000 x g for 10 min and resuspended in 8 µL solubilization buffer A (50 mM NaCl, 50 mM imidazole HCl pH7.0, 2 mM 6-aminohexanoic acid, 1 mM EDTA) by carefully pipetting up and down. To solubilize protein complexes, digitonin was added to obtain a final concentration of 8 g/g and samples were left on ice for 30 min.

Solubilized protein complexes were centrifuged at 20,000 x g for 20 min and supernatants were transferred to LoBind tubes. Glycerol (5% v/v final concentration) and Coomassie blue G-250 (0.6% w/v final concentration) were added and samples were loaded on a 3-12% Bis-Tris NativePAGE gel (Invitrogen). BN-PAGE was performed on ice according to the manufacturer’s instructions. After electrophoresis, the gel was stained using InstantBlue Coomassie protein stain (Abcam).

The BN-PAGE lanes with 0 mM and 0.25 mM DSSO concentrations were each divided into 20 slices. In-gel protein digestion was performed, in short, each slice was transferred into a fresh LoBind tube and destained with a 50% AcN/H2O solution until the Coomassie stain was completely removed. The destained slices were covered with 100% AcN and dehydrated completely. After removing the AcN, enough Dithiothreitol (DTT) was added to cover the gel pieces to achieve a final concentration of 10 mM. The samples were then incubated for 15 minutes at RT. Next, the DTT solution was removed, and the gel pieces were completely dehydrated using 100% AcN. For protein alkylation, the gel slices were covered in a 55 mM Iodoacetic Acid (IAA) solution and incubated in the dark for 30 minutes at RT. The alkylation solution was removed, and the gel pieces were completely dried using 100% AcN. A trypsin buffer was prepared, consisting of 13 ng/µL trypsin (MS Grade, Thermo Scientific™) in 10 mM ammonium bicarbonate (ABC) with 10% (v/v) AcN. The buffer was added to the dried gel slices until each slice was fully covered, and the tubes were incubated at 37°C for 16 hours. The reaction was quenched by acidification with 10% TFA, and the peptides were desalted using C18 StageTips. Subsequently, the peptides were eluted with 80% AcN/0.1% TFA and dried down to prepare for LC-MS/MS analysis.

### LC-MS protein identification of BN-PAGE slices

Peptides from BN-PAGE slices were analysed using an Orbitrap Fusion Lumos Tribrid mass spectrometer (Thermo Fisher Scientific, Germany) connected to an Ultimate 3000 RSLCnano system (Dionex, Thermo Fisher Scientific, Germany), operated under Tune 3.4, SII, and Xcalibur 4.4 software. The mobile phases used were 0.1% (v/v) formic acid (A) and 80% (v/v) acetonitrile with 0.1% (v/v) formic acid (B). Samples were resuspended in 2% acetonitrile with 0.1% formic acid before injection onto an Easy-Spray column (C18, 50 cm, 75 µm ID, 2 µm particle size, 100 Å pore size) operated at 45°C with a flow rate of 300 nl/min. Peptides were eluted using the following gradient: 0 to 2% buffer B in 110 minutes, 2 to 40% B in 1 minute, 40 to 95% B in 11 minutes, and 95 to 2% B in 5 minutes, and the column was then set to washing conditions for 5 minutes at 95% buffer B and recalibrated with 2% buffer B. The mass spectrometer settings were: MS1 resolution of 120,000, AGC target of 100%, maximum injection time set to auto, scan range from 380 to 1200 m/z, and RF lens set at 30%. The DIA settings included a precursor mass range of 400-1200 m/z, quadrupole isolation mode, auto DIA window type, 22 m/z isolation window with 37 scan events, HCD activation type with a normalised collision energy of 33%, Orbitrap detector type with a resolution of 30,000 and a scan range of 300-1600 m/z, RF lens at 30%, custom AGC target with a normalised AGC target of 800%, maximum injection time of 54 ms with custom injection time mode, and positive polarity. Each LC-MS acquisition took 145 minutes.

### MS data analysis of BN-PAGE slices

Raw files obtained from DIA runs from BN-PAGE were processed using DIA-NN 1.8.2 beta 27 (Data-Independent Acquisition by Neural Networks). Output was filtered at a 0.01 FDR threshold, a spectral library was generated based on the provided FASTA file (UniProt Homo sapiens (human) reference proteome (Proteome ID: 9606)), and deep learning was used to create a new in-silico spectral library from the provided peptides. A library-free search was enabled with default settings. Highly heuristic protein grouping was used to reduce the number of protein groups obtained. The legacy (direct) quantification mode was used, normalisation was disabled, and mass accuracy was fixed to 1.5e-05 for both MS2 and MS1. The obtained result file, report.pg_matrix.tsv, was used for complexome profiling of selected proteins, including AIFM1, AK2, and complexes from the OXPHOS. Complexome profiling was performed as follows: for the selected proteins, the intensities were normalised to the highest intensity across the gel fractions. This normalisation was performed separately for both conditions (+/- DSSO).

### Recombinant protein production and purification

E. coli BL21(DE3) (New England Biolabs C2527H) were transformed with 10 ng of the respective expression plasmid and grown on LB agar plates containing 100 µg/mL ampicillin for 18 h at 37°C. Subsequently, a liquid LB overnight culture containing 100 µg/mL ampicillin was inoculated with multiple colonies from the plate and grown under shaking for 18 h at 37°C. 2 L LB medium containing 100 µg/mL Ampicillin was inoculated with 20 mL of overnight culture and grown under shaking in a Multitron HT incubator (Infors) at 37°C, 150 rpm, until OD_600_ = 0.7. Then, the temperature was lowered to 18°C and protein expression was induced by adding 0.2 mM Isopropyl β-d-1-thiogalactopyranoside (IPTG, Carl Roth). Cultures were further grown for 20 h. Bacteria were pelleted by centrifugation at 4000 rpm, 4°C for 15 min using a Fiberlite™ F10-4 × 1000 LEX centrifuge rotor. The pellet was resuspended in 70 mL buffer containing 50 mM Tris-HCl, pH 7.8, 500 mM NaCl, 4 mM MgCl2, 0.5 mM tris-(2-carboxyethyl)-phosphine (TCEP), mini-complete protease inhibitors, EDTA-free (1 tablet per 50 mL, Merck), and 30 mM imidazole. 5 µL Benzonase (Merck) and a spatula tip of lysozyme (Merck) were added, and the suspension was stirred thoroughly. Cells were lysed by sonication using a Sonopuls HD 2070 ultrasound homogeniser (Bandelin) equipped with an MS 72 probe for 3x 5 min at 50% amplitude, 5 s pulse on, and 15 s pulse off. The lysate was clarified by centrifugation at 12000×g for 45 min at 4°C using a Fiberlite™ F15-8 x 50cy centrifuge rotor (Thermo Fisher Scientific), and applied on an XK 16/20 chromatography column (Cytiva) containing 10 mL Ni Sepharose® High-Performance beads (Cytiva), using an Äkta pure FPLC (Cytiva) at 4°C. The column was washed with 250 mL buffer containing 50 mM Tris-HCl, pH 7.8, 500 mM NaCl, 4 mM MgCl2, 0.5 mM TCEP, and 30 mM imidazole. Protein was eluted with the same buffer supplemented with 0.3 M imidazole. Eluent fractions were analysed using SDS-PAGE, and appropriate fractions were pooled and concentrated to 5 mL with centrifugal filter devices (Vivaspin 20, Sartorius). Subsequently, SEC was performed on an Äkta pure FPLC (Cytiva), using a Superdex 200 16/600 column (Cytiva), equilibrated in 10 mM Tris-HCl pH 7.8, 150 mM NaCl, 4 mM MgCl2, 0.5 mM TCEP buffer, at a flow rate of 1 mL/min. SEC fractions were analysed by SDS-PAGE, and appropriate fractions were pooled. The total protein content was estimated by assuming OD_280_ = 1 equals a protein concentration of 1 mg/mL. To cleave off the His-tag, recombinant His-TEV protease was added in a protease:protein ratio of 1:50 and incubated for 18 h on ice. The sample was again applied on an XK 16/20 chromatography column (Cytiva) containing 10 mL Ni Sepharose® High-Performance beads (Cytiva), using an Äkta pure FPLC (Cytiva) at 4°C. The flow-through fraction was collected and concentrated to 5 mL with centrifugal filter devices (Vivaspin 20, Sartorius). A second SEC was performed on an Äkta prime plus FPLC (Cytiva), using a Superdex 200 16/600 column (Cytiva), equilibrated in 10 mM Tris-HCl pH 7.8, 150 mM NaCl, 4 mM MgCl2, 0.5 mM TCEP buffer, at a flow rate of 1 mL/min. SEC fractions were analysed by SDS-PAGE, and appropriate fractions were pooled and concentrated to approx. 20 mg/mL, flash-frozen in liquid nitrogen in small aliquots, and stored at −80°C. Protein concentration was determined with a NanoDrop spectrophotometer (ND 1000, Peqlab), using theoretical absorption coefficients calculated based on the amino acid sequence by ProtParam on the ExPASy web server^83^.

### In vitro reconstitution and analytical SEC

Proteins or protein mixtures were incubated for 1 h on ice at a final concentration of 25 µM per individual protein in a volume of 120 µL, in the presence or absence of 1 mM NADH (Roche). Subsequently, samples were loaded on an analytical SEC column (Superdex 200 Increase 10/300 GL, Cytiva), equilibrated in 10 mM Tris-HCl pH 7.8, 150 mM NaCl, 4 mM MgCl2, and 0.5 mM TCEP, at a flow rate of 0.5 mL/min, using an Äkta pure FPLC (Cytiva). 1 mL fractions were collected and analysed by SDS-PAGE.

### Mass photometry

Mass photometry landing assays were conducted using a Refeyn OneMP instrument (Refeyn Ltd.) with AcquireMP software (version 2023 R1.1). Each condition was measured in triplicate. For the assay, 15 µL of buffer was added to an empty silicon well (CultureWell™ CW-50R-1.0-Gasket, Grace Bio-Labs) placed on a clean coverslip (Menzel-Gläser 24 x 50mm, 1.5 spezial). Then, 5 µL of 20 nM protein stock was added to the well, resulting in a final protein concentration of 5 nM, mixed, and acquisition started immediately. For the measurement of mixed samples of AIFM1 and AK2, equimolar amounts (20 nM) were used. Movies were recorded for 60 seconds at 999 Hz with a frame binning of 10. The system was calibrated with purified Glycinin, which has detectable oligomeric species at molecular weights of 160 kDa, 320 kDa, 480 kDa, and 640 kDa, to convert contrast to mass. Data was processed using Discover MP software (version 2023 R1.2) to obtain mean peak contrast and mass values through Gaussian fitting.

### Cryo-EM sample preparation

The AIFM1/AK2 protein complex for cryo-EM was prepared as described above (*in vitro* reconstitution and analytical SEC). R1.2/1.3 400 mesh Cu holey carbon grids (Quantifoil) were glow-discharged for 30 s using a Fischione Model 1020 plasma cleaner and a 95% argon and 5% oxygen mixture. 3.5 μl solution containing 0.5 μM (0.07 mg/mL) of pooled SEC peak fractions corresponding to the AIFM1/AK2 complex were applied to the grid, incubated for 45 s, blotted with a Vitrobot Mark IV device (ThermoFisher) for 1–2 s at 4°C and 99% humidity and plunged in liquid ethane. Grids were stored in liquid nitrogen until imaging.

### Cryo-electron microscopy and data processing

The data set was collected on a Titan Krios G3i transmission electron microscope (ThermoFisher Scientific, Server version 2.15.3, TIA version 5.0) operated at an acceleration voltage of 300 kV and equipped with an extra-bright field emission gun, a BioQuantum post-column energy filter (Gatan), and a K3 direct electron detector (Gatan, Digital Micrograph version 3.32.2403.0). All images were acquired in low-dose mode as dose-fractionated movies using EPU version 2.8.1 (ThermoFisher) with a maximum image shift of 12 µm using aberration-free image shifting.

2535 movies with a total dose of 60 e Å^−2^ each divided into 60 fractions were collected in energy-filtered zero-loss (slit width 20 eV) nanoprobe mode at a nominal magnification of 105.000x (resulting in a calibrated pixel size of 0.425 Å at the specimen level) in super-resolution mode. Data were recorded with defocus values ranging from −0.5 to −2.5 µm.

MotionCor2^84^ was used for motion correction. Initial estimation of the contrast transfer function (CTF) was performed with the CTFFind4 package^85^ via the Relion 3.0.7 GUI^86^. Power spectra were manually inspected to remove ice contamination and astigmatic, weak or poorly defined spectra. Particles were picked using a blob picker via the Relion 3.0.7 GUI, resulting in 1,748,043 particles. Decimated particles were extracted at an initial pixel size of 3.4 Å/pixel (Fig. S4A). All subsequent steps were performed in CryoSPARC 4.2.1^52^. Particles were subjected to one round of multi-class ab initio reconstruction (4 classes, class similarity = 0), resulting in one defined AIFM1 dimer class. This class exhibited highly fragmented density corresponding to AK2. Subsequently, all particles were reassigned using the maps derived from the ab initio reconstruction, resulting in 794,854 particles assigned to the AIFM1 dimer class. All AIFM1 particles were refined using homogeneous refinement. It was noted that the auto-masking resulted in mask artefacts in flexible regions. Therefore, the auto-masking option was disabled during the refinement. AIFM1 dimer particles were re-extracted at the full pixel size (0.85 Å/pixel) and underwent one round of 3D variability analysis using a filter resolution of 5 Å. The variability display indicated that AK2 was either partially absent, fragmented, or associated with the AIFM1 dimer in varying positions. Six clusters were chosen, which resulted in four cluster projections with defined AK2 densities, one with fragmented density, and one where no AK2 could be observed. Subsequently, particles were reassigned to three of the cluster projections (“no AK2”, “AK2 leaned forward”, “AK2 leaned back”). Further heterogeneity was identified using hierarchical multi-class ab initio reconstructions. Identified subclasses were used for another round of heterogeneous refinement. Finally, classes with the most stable conformations (“AK2 leaned forward” (class 3): 4.3 Å, “intermediate AK2 position” (class 2): 6.2 Å and “AK2 leaned back” (class 1): 4.6 Å.) of AK2 were subjected to non-uniform refinement.

### Model building and refinement

The best-resolved map (class 3) was used for rigid body fitting of the crystal structure of dimeric reduced AIFM1 (PDB ID 4bur^50^) using the program ChimeraX 1.7.1^87^. Coordinates corresponding to AK2 and the AIFM1 C-domain containing the AK2 C-terminal insertion were taken from the AlphaFold prediction and superposed with the 4bur C-domain using ChimeraX 1.7.1. The model was adjusted manually using the program Coot 0.9.8.7^88^, and automated real-space refinement was carried out using the program Phenix 1.20.1-4487^89^.

### Molecular dynamics simulations

All the MD simulations were performed using GROMACS 2020.4 software^90^ under the effect of CHARMM36 force-field^91^. The simulation box was chosen as a dodecahedron and the AIFM1-AK2 complex was located 14 Å from the edges of the box and solvated with TIP3P^92^ water molecules. Then the system was neutralized with 0.15 M with Na+ and Cl-ions, followed by the system minimization with steepest descent algorithm. After energy minimization, the system was heated to 300 K under constant volume for 100 picoseconds (ps) long. During this period, positional restraints were applied on heavy atoms with 1000 kJ/mol.nm2 force constant. While keeping the number of particles in the system constant, by using the Parrinello-Rahman barostat^93^, the NPT equilibration of the system was enforced in 100 ps. The MD simulations were run under zero positional restraints for 2 µs long. Two replica simulations were carried out for each system (#1 2x(AIFM1(NAD+FAD):AK2, 2×2µs), #2 2x(AIFM1(FAD):AK2, 2×2µs)) starting with different initial velocities (i.e., random seeds). During the simulation, the PME approach^94^ was employed to address electrostatic interactions, utilizing a nonbonded cutoff of 1.2 nm. A fourth-order LINCS^95^ method was used to restrict bond lengths. The time step was set to 2 fs in every simulation. Before the analysis part, the equilibration time was chosen as 300 ns for AIFM1(FAD):AK2 and 500 ns for AIFM1(NAD+FAD):AK2. These time periods were discarded during the analysis, leading practically to 2×1.7µs for AIFM1(NAD+FAD):AK2 and to 2×1.5µs production run times (see Fig. S5A, B).

### AP-MS

For transfection, 293T cells were seeded in triplicate 145 mm plates (Greiner Bio-One). Ten million cells in 15 mL DMEM (Corning cat# 15-013-CV) supplemented with 10% FBS (Sigma-Aldrich cat# S0615-500ML) and 2 mM L-glutamine (Corning cat# 25-005-CI) were seeded per plate and incubated for 20 h. Cells were washed with PBS (Corning cat# 21-031-CV) and transfected with 15 µg pcDNA3.1-FLAG-AK2 plasmid per plate using the PolyJet DNA transfection reagent (SignaGen), according to the manufacturer’s instructions, and incubated for 44 h. If applicable, sodium iodoacetate (Sigma-Aldrich, cat# 57858-5G-F) was added from a 0.5 M stock in PBS to a final concentration of 100 µM, and the plates were incubated for a further 4 h. Cells were washed with PBS, detached with trypsin/EDTA solution, counted, and washed again with PBS. If applicable, the cells were then resuspended in PBS containing 6 mM SDA (added from a 0.2 M stock in DMF) for crosslinking at a cell density of 28×10^6^ cells/mL. Cells were rotated at 22°C for 30 min. Tris-HCl (Santa Cruz Biotechnology) was added to a final concentration of 50 mM, cells were rotated for an additional ten minutes at 22°C, and washed with PBS. Subsequently, cells were irradiated with UV light at 365 nm using a Luxigen LZ1 LED emitter (Osram Sylvania Inc.) to form crosslinks for 2×10 s at a cell density of 28×10^6^, and washed with PBS. All cell pellets were flash-frozen in liquid nitrogen and stored at −20°C.

Cell pellets were resuspended in 1 mL lysis buffer per 40×10^6^ cells, containing 42 mM Na_2_HPO_4_ (AppliChem), 8 mM NaH_2_PO_4_ (AppliChem),500 mM NaCl (Carl Roth), 10% glycerol (AppliChem), 4 mM MgCl2 (Carl Roth), 0.5 mM TCEP (Carl Roth), 0.5% LMNG, 0.005% CHS (Anatrace cat# NG310-CH210), protease inhibitors (mini complete EDTA-free, Roche, 1 tablet per 50 mL), 125 U benzonase (Millipore cat# E1014). The suspension was passed 5x through a 26G syringe needle and rotated for 20 min at 4°C. The lysate was clarified by centrifugation at 10,000 and 4°C for 20 min. 50 µL of washed anti-FLAG affinity beads (Sigma-Aldrich cat# A2220) were added to 1 mL clarified lysate and rotated for 2.5 h at 4°C. Subsequently, the beads were washed with 1 mL lysis buffer, then twice with 1 mL lysis buffer without LMNG/CHS, and once with 1 mL 10 mM HEPES pH 7.5, 10% glycerol. Afterwards, 10 µL of the beads were removed for western blotting, and the remaining 40 µL proceeded to on-bead proteolytic digestion. To this end, the beads were resuspended in 50 µL buffer containing 50 mM (NH_4_)HCO_3_ (AppliChem), 6 M urea (AppliChem), 2 M thiourea (Sigma-Aldrich). DTT was added from a fresh 1 M stock in 50 mM (NH_4_)HCO_3_ to a final concentration of 20 mM, and the bead suspension was incubated under shaking using an Eppendorf ThermoMixer at 1000 rpm, 25°C, for 15 min. Subsequently, iodoacetamide was added from a 1 M stock in 50 mM (NH_4_)HCO_3_ to a final concentration of 125 mM and incubated for a further 30 min under shaking at 1000 rpm, 25°C. The suspension was diluted with 300 µL (NH_4_)HCO_3_, 0.5 µg of trypsin (Thermo Scientific cat# 90057) was added from a 1 µg/mL stock solution in 0.1% TFA, and incubated for 20 h under shaking at 1000 rpm, 25°C. The resulting tryptic peptides were desalted using C18 StageTips^81^ and fractionated on a Superdex Peptide 3.2/300 increase column (GE Healthcare) at a flow rate of 10 μl/min using 30% (v/v) acetonitrile and 0.1% (v/v) trifluoroacetic acid as mobile phase. 50 μl fractions were collected, and peptide-containing fractions were pooled and vacuum-dried. Peptide concentrations were estimated by integrating the 214 nm UV-absorption peak area, and equal amounts of peptides were resuspended for mass spectrometry as detailed below.

For the FLAG-AK2 pulldowns ∓ in-cell SDA crosslinking and the corresponding empty vector controls, an amount corresponding to 8 mAU_214_•mL per sample of vacuum-dried peptides from pooled SEC fractions was resuspended in 250 µL 0.1% (v/v) formic acid, and 10 µL were loaded on the Evotip™ using the standard producer protocol in quadruplicates. We utilized EVOSEP coupled with a TimsTOF ULTRA (Bruker) mass spectrometer equipped with a Captive Spray II source (Bruker). The separation was carried out on a Performance Column OE measuring 8 cm x 150 µm ID, with a particle size of 1.5 µm, maintained at 40°C. We employed the standard 60SPD EVOSEP method, applying a 21-minute gradient for a total sample-to-sample time of 24 minutes using Solvent A: 0.1% (v/v) formic acid (FA) and Solvent B: acetonitrile (ACN)/0.1% (v/v) FA (Thermo Fisher Scientific™ Optima LCMS grade). The dia-PASEF acquisition scheme was optimized for a cycle time estimate of 1.20 s. The window scheme was designed to cover most of the charge 2 precursor ions in the range m/z 392−1113 and 1/K0 0.67−1.34, using 26 × 28.7 Th windows, with accumulation and ramp times of 85 ms. The mass spectrometer was operated in the “high sensitivity detection” (“low sample amount”) mode.

For the samples arising from iodoacetic acid treatments and corresponding controls, LC-MS/MS analysis was carried out using an Orbitrap Fusion Lumos Tribrid mass spectrometer (Thermo Fisher Scientific) coupled online with an Ultimate 3000 RSLCnano system (Dionex, Thermo Fisher Scientific). An amount corresponding to 6 mAU_214_•mL per sample of vacuum-dried peptides from pooled SEC fractions was resuspended in 12 µL 0.1% (v/v) formic acid, 1.6% (v/v) acetonitrile. Triplicate 3 µL injections of each sample were performed. The peptides were separated on a 50 cm EASY-Spray C18 column (Thermo Scientific) operating at 45°C column temperature. Mobile phase A consisted of 0.1% (v/v) formic acid and mobile phase B of 80% v/v acetonitrile with 0.1% v/v formic acid. Peptides were loaded at a flow rate of 300 nL min−1 and separated at a flow rate of 250 nL min−1. Peptides were separated using a gradient with linear increases from 2% mobile phase B to 35% over 65 minutes then to 45% over 7 minutes, followed by a steep increase to 90% mobile phase B in 5 min.

Eluted peptides were ionized by an EASY-Spray source (Thermo Scientific) and introduced directly into the mass spectrometer. The MS data were acquired in the data-independent (DIA) mode. For every acquisition cycle, one MS1 spectrum was recorded in the Orbitrap at a resolution of 60K with an m/z range of 390-1100. Subsequently, 38 DIA MS2 scans were acquired with an isolation m/z window of 16 (0 overlap) and a precursor m/z range of 400-1000. Precursors were fragmented employing higher-energy collisional dissociation (HCD) with a collision energy of 33%. DIA spectra were recorded in the Orbitrap at a resolution of 30K, the scan range was 300-1600 and the maximum injection time was set to 54 ms.

Raw files obtained from DIA runs were processed using DIA-NN 1.8.2 beta 27. A modified FASTA file (UniProt Homo sapiens (human) reference proteome (Proteome ID: 9606)) was provided, where the FLAG affinity tag amino acid sequence was inserted in the corresponding AK2 FASTA entry. A library-free search was enabled with Trypsin/P protease, 2 missed cleavages, peptide length range 7-30, precursor charge range 1-4, precursor m/z range 300-1800, fragment ion m/z range 200-1800. Cysteine carbamidomethylation was set as a fixed modification. Precursor FDR was set to 1%, and MBR was activated. All other settings were left as default. Using the resulting “report.tsv” file as input, protein enrichment analysis of FLAG-AK2-transfected cells, compared to empty vector controls, in the presence and absence of in-cell SDA crosslinking (Fig. 1C) was conducted using DEqMS differential abundance analysis^96^ as implemented in the MS-DAP platform^47^, with the following settings: min_detect = 2; fraction_detect = 0.66; min_quant = 2; fraction_quant = 0.66; norm_algorithm = ‘vsn&modebetween_protein’; rollup_algorithm = ‘maxlfq’. For quantification of the AIFM1/AK2 peptide intensity ratio (Fig. 5B), the “report_pr_matrix.tsv” output from DIA-NN analysis was used. AIFM1 and AK2 peptides were identified which were quantified across all biological and technical replicates, and their intensities were summed up. Subsequently, the sum of AIFM1 intensities was divided by the sum of AK2 intensities. Statistical analysis of the resulting ratios was done by a two-sample t-test.

### Immunoblotting

For western blotting, 5 µL of the cleared lysates and 10 µL of washed beads after pull-down were applied on separate Mini-PROTEAN TGX 4-12% polyacrylamide gels (BIO-RAD), and run for 45 min at 200 V in Tris/Glycine/SDS buffer. Separated proteins were transferred on PVDF membranes (Trans-Blot Turbo Transfer Pack, BIO-RAD) using a Trans-Blot Turbo Transfer System (BIO-RAD) according to the manufacturer’s instructions. Membranes were cut horizontally above the 38 kDa marker band and blocked in 5% milk powder in TBS-T for 2 h at 22°C under slight agitation. The upper parts of the membranes were then incubated for 20 h at 4°C with mouse monoclonal anti-AIFM1 antibody (Aviva cat# OAAH00022) at a dilution of 1:3500 in 5% milk powder in TBS-T, and the lower parts were incubated or 20 h at 4°C with mouse monoclonal anti-FLAG antibody (Sigma-Aldrich cat# F1804) at a dilution of 1:5000 in 5% milk powder in TBS-T, under slight agitation. The membranes were washed three times for 5 min at 22°C with TBS-T, and subsequently incubated for 2.5 h at 22°C with goat anti-mouse-HRP secondary antibody (Sigma-Aldrich cat# A3682) at a dilution of 1:10000 in 5% milk powder in TBS-T, under slight agitation. The membranes were washed three times for 5 min at 22°C with TBS-T, under slight agitation. Blots were developed using SuperSignal West Pico PLUS chemiluminescent substrate (ThermoFisher) in a ChemiDoc XRS+ chemiluminescence reader (BIO-RAD), using the Image Lab software (BIO-RAD).

### C. elegans methods

The *let-754* allele without exon-5 was engineered using CRISPR/Cas9-mediated genome editing using the Cas9/RNP method^97^. To generate the *let-754 Δ*exon-5 allele a single-stranded repair template with a 35 bp homology arm was used. Gene editing was performed by injecting the repair template and a ribonucleoprotein (RNP) mixture comprising crRNA/tracrRNA (Integrated DNA Technologies) and purified recombinant Cas9 enzyme into the gonads of young N2 adults. The injection mix also contained RNPs targeting the R92C mutation in dpy-10, which results in a dominant roller phenotype^98^, to select for animals with potentially successful edits. The roller worms were further screened by genotyping PCRs using primers that track the desired genome deletion (*let-754 Δ*exon-5). Positive clones from the PCR were then confirmed by sequencing. The CRISPR strain was outcrossed multiple times to remove any background and off-target mutations.

For the brood size assay, all animals were reared up to the L4 stage at 20°C on 55 mm unvented Petri plates with 1x NGM (Nematode Growth Medium), seeded with 200 µL E. coli OP50 bacteria. Plates were seeded with OP50 no longer than one week before use. Rearing plates were wrapped in Parafilm during storage, but experimental plates were not. On day 0, individual L4 animals from the rearing plate were transferred onto fresh experimental OP50 plates (Day 1 plates, one animal per plate) and stored upside down at 20°C or 25°C. Animals were transferred onto new experimental plates every 24 h until day 5, then terminated. Experimental plates were allowed to develop at their respective temperatures for another approximately 24 h after the removal of the egg-laying animal until progeny reached the L4 stage. At this point, all living offspring were counted and removed from the plate. Unhatched eggs were then counted separately and not included in the brood size numbers. Brood size was calculated by summing living offspring counts from experimental plates corresponding to days 1-5 for each animal in the assay. Statistical analysis was done using the Wilcoxon matched-pairs signed rank test., with a p-value of 0.0078 for the 20°C data set and 0.0098 for the 25°C dataset.

For the lifespan assay, 20-25 *C. elegans* animals at larval stage L4 were picked onto a 55 mm diameter 1x NGM growth plate seeded with 200 μL *E. coli* OP50 bacteria (two plates per strain) and stored at 20°C in the dark for approximately 72 h. From the resulting progeny, 15 animals at stage L4 were picked onto a 55 mm diameter 1x NGM assay plate seeded with 200 μL OP50. Ten assay plates were used per strain for a total of n = 150 animals per strain. Picking day was designated Day 0. The assay plates were stored in a plastic box lined with aluminium foil to minimize light exposure and kept at 28°C for the duration of the assay. Animals were transferred onto fresh OP50 assay plates on days 2 and 5 only to minimize handling. Surviving animals were counted daily. In the case of old, non-mobile animals, the following sequence was used to assess whether the animals were still alive by evaluating responses to mechanical stimulation: (1) tapping the plate 2-3 times; (2) gentle touch with pick on tail of animal; (3) gentle touch with pick on body of animal, close to middle; (4) gentle touch with pick on head of animal. Steps in the sequence were performed approximately 15 s apart. Animals meeting one of the exclusion criteria (escaped from plate/not found; dried on plate wall; vulva protrusion; vulva rupture; paralysis; bag of worms; death due to mishandling by experimenter) were censored from the assay. Resulting data were plotted as Kaplan-Meier curves and statistical comparison was performed using the log-rank (Mantel-Cox) test in GraphPad Prism 9. The experiment was conducted in triplicate.

For the L1 recovery assay, synchronized L1 arrested animals suspended in M9 were divided into two sets of 12 culture tubes under sterile conditions, ensuring equal numbers in each tube. These sets were then cultured at 20°C and 25°C, shaking at 30rpm for 24 days. Every two days, triplicate 10 µL droplets from the same culture tubes at either 20°C or 25°C were transferred onto NGM plates seeded with OP-50 E. coli and incubated at 20°C for 48 hours. After two days, the total number of animals that developed to different stages was calculated. The percentage of L4 animals per day was calculated as 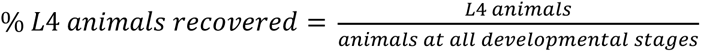.

## Data availability

MS data have been deposited at ProteomeXchange via the PRIDE partner repository with the dataset identifiers PXD052832 (crosslinking mass spectrometry) (Username; Password: jIQbPdhNDTUh) and PXD052817 (linear protein identification of soluble and insoluble fraction) (Username; Password: 3IDoDH6AyssY), PXD052821 (complexome profiling) (Username; Password: AbJ7IqCqkIGi), PXD052867 (FLAG-AK2 *∓*SDA pull-downs) (Username; Password: fotKriDBeqKn) and PXD052852 (FLAG-AK2 *∓* IA pull-downs) (Username; Password: 9WD6wGq4A7qq).

The EM maps have been deposited at the Electron Microscopy Data Bank (EMDB) under the accession codes EMD-50534 (class 1), EMD-50535 (class 2), and EMD-50532 (class 3). Atom coordinates have been deposited at the Protein Data Bank (PDB) with the accession number 9FL7.

MD trajectories will be uploaded to Zenodo before publication.

## Acknowledgements

We thank Michelle Krause for her help with the analytical SEC, as well as Anne Schulze and Dr. Lukasz Szyrwiel for assistance with mass spectrometry. We thank the Core Facility for cryo-EM (CFcryoEM) of the Charité - Universitätsmedizin Berlin for support in the acquisition of the data. The CFcryoEM was supported by the German Research Foundation (DFG) through grant No. INST 335/588-1 FUGG. The work was supported by the DFG under Germany’s Excellence Strategy—EXC 2008—390540038—UniSysCat (JR, FS, SML and CMTS), project 329673113 (JR), and the Emmy Noether programme (SCHW1851/1-1) (DS). DC was supported by the Wellcome Trust and the Royal Society (208833/Z/17/Z). SML and CMTS were supported by the Bundesministerium für Bildung und Forschung (BMBF 16GW0300 to CMTS). The Wellcome Centre for Cell Biology is supported by core funding from the Wellcome Trust (203149). The Genotype-Tissue Expression (GTEx) Project was supported by the Common Fund of the Office of the Director of the National Institutes of Health, and by NCI, NHGRI, NHLBI, NIDA, NIMH, and NINDS. The data used for the analyses described in this manuscript were obtained from the GTEx Portal Transcript Browser using the search term “AK2” on 14 May 2024.

## Author contributions

All authors planned experiments

FS, PSJR, ABB, VRO, KJ, DM, MS, KS, DS performed experiments

FS, PSJR, SML, SL, ABB, VRO, KJ, DM, FJO’R, AG, KS, EK, KEB, DC, DS, JR analysed data

HE, CMTS, EK, KEB, DC, DS, JR supervised the research

DS, JR drafted the manuscript, all authors edited and approved the manuscript

## Competing interests

The authors declare no competing interests.

---

**Table S1:**
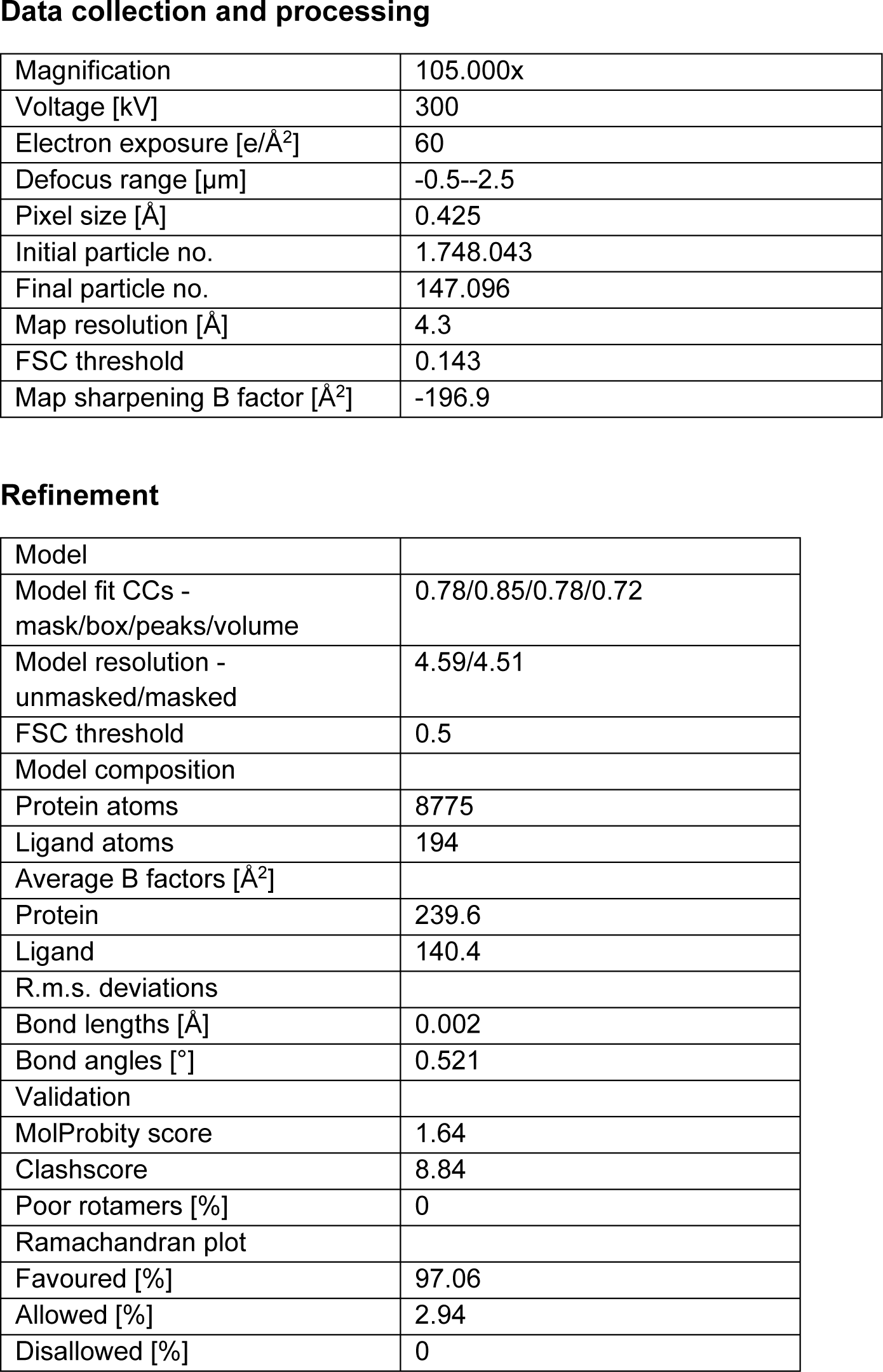
Cryo-EM data collection and refinement Data collection and processing.

|  |  |
| --- | --- |
| Magnification | 105.000x |
| Voltage [kV] | 300 |
| Electron exposure [e/Å<sup>2</sup>] | 60 |
| Defocus range [μm] | -0.5--2.5 |
| Pixel size [Å] | 0.425 |
| Initial particle no. | 1.748.043 |
| Final particle no. | 147.096 |
| Map resolution [Å] | 4.3 |
| FSC threshold | 0.143 |
| Map sharpening B factor [Å<sup>2</sup>] | -196.9 |
| **Refinement** |  |
| Model |  |
| Model fit CCs - mask/box/peaks/volume | 0.78/0.85/0.78/0.72 |
| Model resolution - unmasked/masked | 4.59/4.51 |
| FSC threshold | 0.5 |
| Model composition |  |
| Protein atoms | 8775 |
| Ligand atoms | 194 |
| Average B factors [Å<sup>2</sup>] |  |
| Protein | 239.6 |
| Ligand | 140.4 |
| R.m.s. deviations |  |
| Bond lengths [Å] | 0.002 |
| Bond angles [°] | 0.521 |
| Validation |  |
| MolProbity score | 1.64 |
| Clashscore | 8.84 |
| Poor rotamers [%] | 0 |
| Ramachandran plot |  |
| Favoured [%] | 97.06 |
| Allowed [%] | 2.94 |
| Disallowed [%] | 0 |

## Notes

### Competing Interest Statement

The authors have declared no competing interest.

